# The 2023 wildfire season in Québec: an overview of extreme conditions, impacts, lessons learned and considerations for the future

**DOI:** 10.1101/2024.02.20.581257

**Authors:** Yan Boulanger, Dominique Arseneault, Annie Claude Bélisle, Yves Bergeron, Jonathan Boucher, Yan Boucher, Victor Danneyrolles, Sandy Erni, Philippe Gachon, Martin P. Girardin, Eliane Grant, Pierre Grondin, Jean-Pierre Jetté, Guillemette Labadie, Mathieu Leblond, Alain Leduc, Jesus Pascual Puigdevall, Martin-Hugues St-Laurent, Junior A. Tremblay, Kaysandra Waldron

## Abstract

The 2023 wildfire season in Québec set records due to extreme warm and dry conditions, burning 4.5 million hectares and indicating persistent and escalating impacts associated with climate change. The study reviews the unusual weather conditions that led to the fires, discussing their extensive impacts on the forest sector, fire management, boreal caribou habitats, and particularly the profound effects on First Nation communities. The wildfires led to significant declines in forest productivity and timber supply, overwhelming fire management resources, and necessitating widespread evacuations. First Nation territories were dramatically altered, facing severe air quality issues and disruptions. While caribou impacts were modest across the province, the broader ecological, economical, and social repercussions were considerable. To mitigate future extreme wildfire seasons, the study suggests changes in forest management practices to increase forest resilience and resistance, adapting industrial structures to new timber supplies, and enhancing fire suppression and risk management strategies. It calls for a comprehensive, unified approach to risk management that incorporates the lessons from the 2023 fire season and accounts for ongoing climate change. The study underscores the urgent need for detailed planning and proactive measures to reduce the growing risks and impacts of wildfires in a changing climate.

## 1. Introduction

Fueled by record-breaking warm and dry conditions (ESCER 2023; Barnes et al. 2023), the 2023 fire season that occurred in Québec was one of extremes. By the end of October, more than 4.5 Mha of forest had burned throughout the province (SOPFEU 2023a), doubling the previous record set in 1989 (2.3 Mha). The total area burned within the commercial forest, which also corresponds to the Intensive Protection Zone under fire management, reached approximately 1.1 Mha, the highest since 1923. These numbers are roughly equal to the area that has burned in the Intensive Protection Zone over the past 20 years combined. Extreme conditions for fire spread fueled the largest wildfire events ever observed in both the Intensive Protection Zone (460 kha) and the northern protection zone where fire management is extensive (1 Mha) (CWFIS 2023).

On June 1st, due to the numerous and fast spreading wildfires, the provincial authorities responsible for fire management in Québec declared that they had reached preparedness level 5, (CIFFC 2023), meaning that this situation had created extreme demands for provincial firefighting resources. Such a preparedness level resulted in the mobilization of national and international resources to assist the *Société de protection des forêts contre le feu* (SOPFEU), the agency responsible for fire management and suppression (CIFFC 2023). The situation was particularly hazardous as dangerous fires were burning near communities, leading to a significant number of fire-related evacuations, most notably by First Nation communities (Canadian Forest Service 2023). Additionally, the air quality was severely compromised, threatening the health of a large proportion of the population of the province, living up to several hundreds of kilometers away from the blazes (CBS News 2023). The dense smoke plumes originating from the 2023 fires in Québec also led to air quality alerts on multiple occasions for large areas in adjacent provinces as well as in northern United States (The New York Times 2023). Eventually, the smoke plumes would cross the Atlantic and cause hazy skies in western Europe (Leibniz Institute for Tropospheric Research 2023).

The 2023 fire season in Québec was a shock for many, as the previous decade had been particularly quiet in terms of wildfires (CIFFC 2023). However, several studies had warned of increased fire activity as a result of increased anthropogenic climate forcing, with potential dire consequences, notably for Québec’s forest sector (Bergeron et al. 2010; Gauthier et al. 2015; Chaste et al. 2019) as well as on infrastructures and communities (Erni et al. 2021). Although consequences of these fires were immediate for communities, public safety, and industries, such an extensive and record-breaking fire season will no doubt have longer-term impacts on several important aspects of Québec’s forest ecosystems, economy, and society. In the following sections, we report to which point this season was exceptional in the context of historical variability. Furthermore, we summarize the impacts the 2023 wildfire season had and will have on i) the forest sector, ii) fire management, communities, and infrastructures, iii) wildlife and their habitats and iv) First Nations in the province. We also provide potential avenues to mitigate these impacts.

## 2. The 2023 fire season in Québec: a timeline and a historical perspective

### 2.1. The 2023 weather conditions that fueled this extreme fire season

The province of Québec, with over 4.5 million hectares burned in 2023, was Canada’s most affected province by wildfires. Most fires occurred in the western and northwestern parts of the province, specifically within the Eastern James Bay and Eastern Subarctic Homogeneous Fire Regime (HFR) zones, with 12% (the highest annual rate of any HFR in Canada since at least 1980) and 2.5% of their areas burned, respectively (Figure 1).

**Figure 1.**
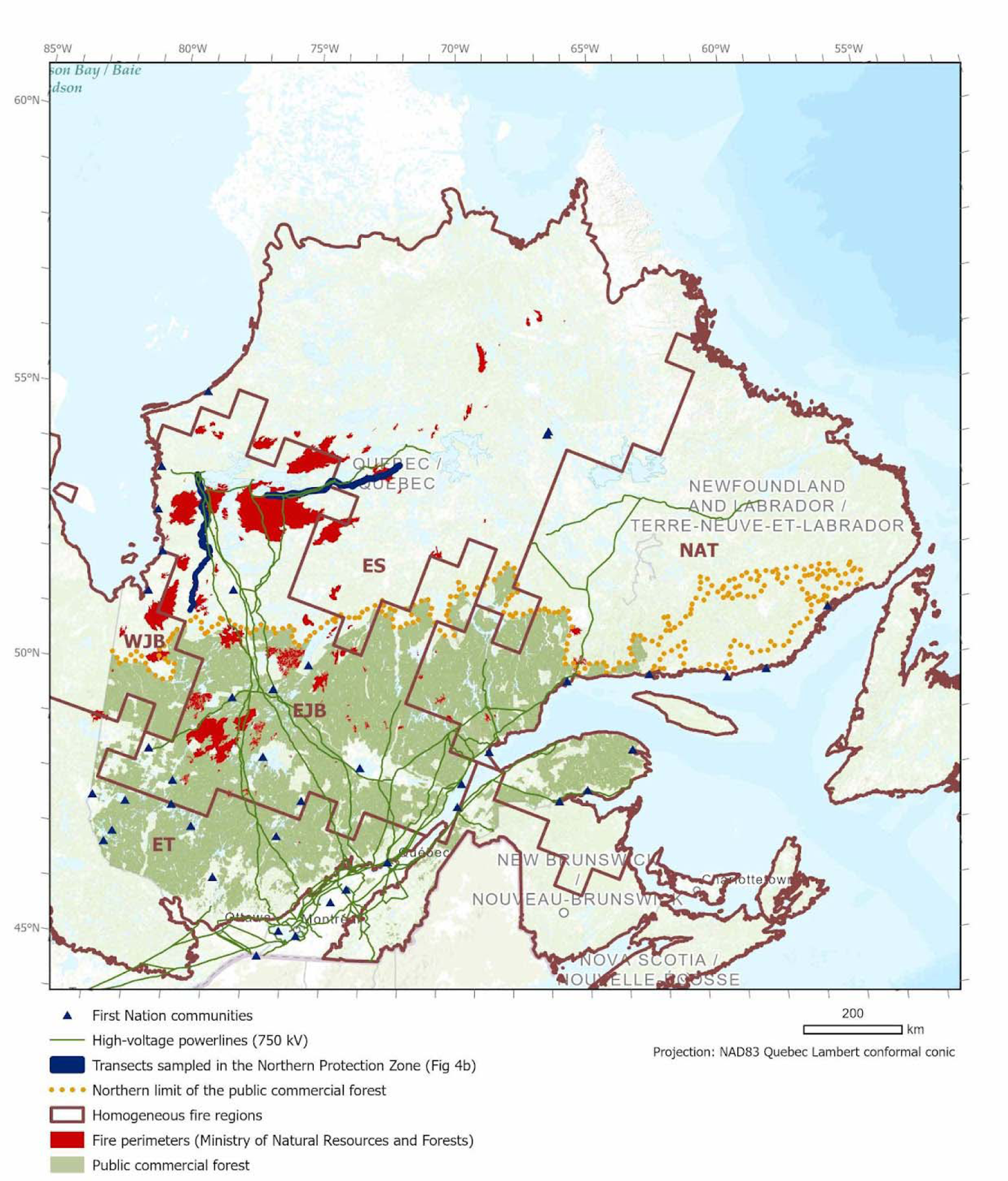
Location of the 2023 wildfires in Québec (April 14th to October 1st) as mapped by the *Direction de la protection des forêts* of Québec’s *Ministère des Ressources Naturelles et des Forêts*. Only fires above 1,000 ha are mapped. The public commercial forest is delineated according to public forest management units and the northern forest allocation limit. The Intensive Protection Zone is located south of this limit whereas the northern protection zone is to its north. Homogeneous fire regime (HFR) zones were retrieved from Boulanger et al. (2014). EJB: Eastern James Bay HFR zone; ES: Eastern Subarctic HFR zone; NAT: North Atlantic HFR zone; ET: Eastern Temperate HFR zone; WJB: Western James Bay HFR zone. High-voltage powerline data are retrieved from *Ministère des Ressources naturelles du Québec* (2019). Transects sampled for fire history in the James Bay area (see section 2.2) are also shown.

Québec experienced its warmest May to August period since 1950, with maximum daily temperatures reaching record highs (Figure 2). Although overall precipitation was near normal, the Eastern and Western James Bay HFR zones had one of their driest seasons on record, with the areas along the James Bay area experiencing Canada’s driest anomalies (Suppl. Mat. S1.4). June saw the highest precipitation deficits, especially in the northwestern and northern parts of the province, persisting for most of the summer along the James Bay coast (Suppl. Mat. S1.1). Temperatures from April to July were consistently above average across the province, peaking in June with areas in the northern half of the province experiencing anomalies of over +3°C, up to +5°C (Suppl. Mat. S1.1). An early snowpack melt in April (Suppl. Mat. S1.2), leading to the lowest May snow water equivalent since 1950 (Figure 2), facilitated the early onset and intensity of the wildfire season. Fire-Weather Index (FWI) for May through August reached unprecedented levels since 1950, significantly above the 1991-2020 normal (Figure 2). The Eastern James Bay and Eastern Subarctic HFR zones saw the highest FWI anomalies, especially in June and July (Suppl. Mat. S1.1), the period during which most fires burned.

**Figure 2.**
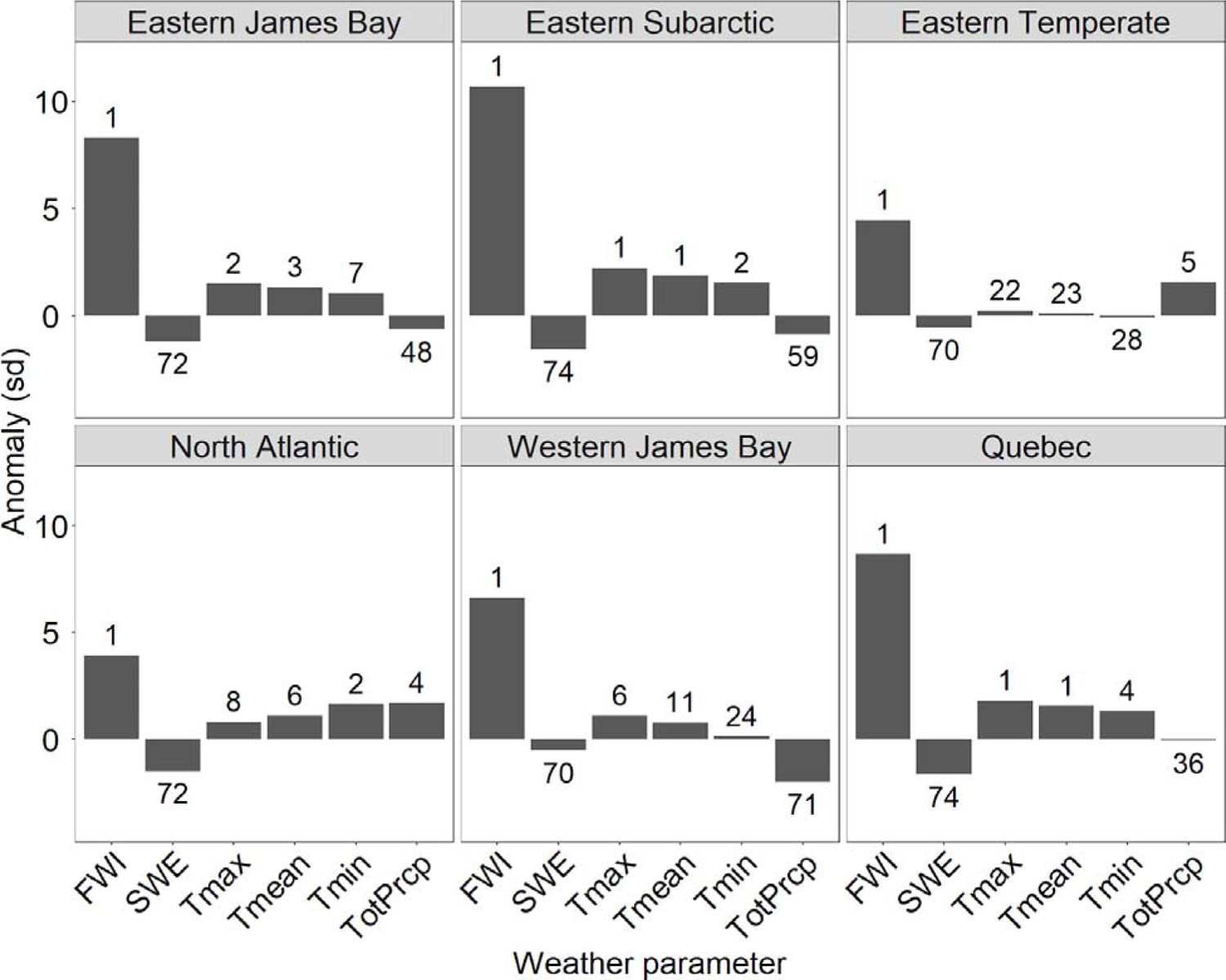
Anomalies (expressed as standard deviations) for various weather parameters for the May-August 2023 period compared with the 1991-2020 normals for each homogeneous fire regime zone (see Figure 1), as well as for all zones combined (Québec). Ranking of 2023 values (from 1: highest to 74: lowest) against the 1950-2023 period is also shown for each parameter. FWI: Fire-weather index; TotPrcp: Total precipitation; Tmax: Maximum daily temperature; Tmean: Mean daily temperature; Tmin: Minimum daily temperature; SWE: snow water equivalent. Data were calculated from ERA5 reanalyses (Hersbach et al. 2020).

Overall, these conditions fueled an intense wildfire season that lasted from late May through September, peaking during three intervals (June 1-12, June 19-28, July 3-15) where daily burned areas often exceeded 100,000 ha, coinciding with sustained high to extreme FWI values (Figure 2). Of note, early June thunderstorms triggered over 120 wildfires, rapidly spreading due to high FWI values (Figure 3). Persistent dry conditions from June to mid-July (Figure 2) supported ongoing and new fires in the Eastern and Western James Bay and Eastern Subarctic HFR zones. Fire progression decreased after mid-July as FWI values dropped in the Eastern James Bay HFR zone (Figure 3), where most active fires were located.

**Figure 3.**
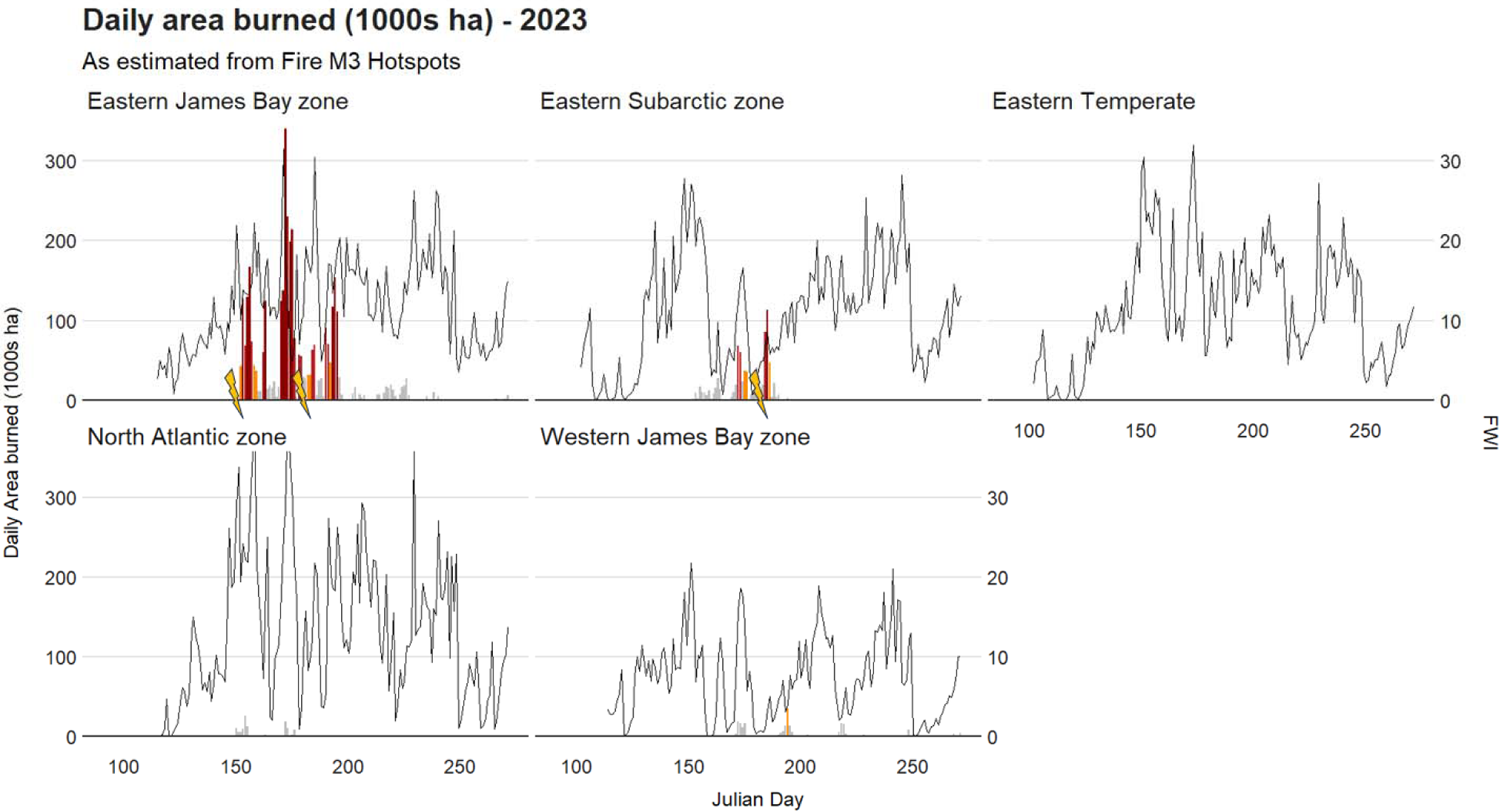
Daily area burned (in thousands of ha, gray bars: less than 30 kha burned; Orange bars: 30 - 50 kha burned; Red bars: 50 - 100 kha burned; Dark red bars >100 kha burned) within each homogeneous fire regime overlapping Québec during the 2023 fire season. Daily fire-weather indices spatially averaged for the whole HFR zone and as assessed from ERA5 reanalyses (Hersbach et al. 2020, black lines) are also shown for the same period. Lightnings correspond to periods where large numbers of lightning-caused fires were ignited.

### 2.2. The 2023 fire season relative to the historical range of variability

The comparison of weather conditions and fire activity in 2023 with those of the past is essential to better understand their exceptionality in the context of climate change. An analysis by the World Weather Attribution initiative (Barnes et al. 2023) concluded that both the intensity (FWI values averaged over a 7-day period) and the severity (cumulative daily severity rating, Van Wagner 1987) of weather conditions from the beginning of the fire season through the end of July 2023 were respectively at least twice and seven times more likely because of the ongoing anthropogenic climate change. These results are in line with several other analyses that have concluded that many wildfire-weather metrics are being modified by climate change in Québec including lengthier fire seasons (Jain et al. 2017), more severe and frequent extreme fire-weather indices (Jain et al. 2022), and drier fuels (vegetation) (Ellis et al. 2022). Most of these changes are actually occurring in northwestern and northern Québec, the very region where the 2023 fire activity was concentrated.

The Drought Code (DC) records of 2023 showed that, as of May 1st in northern Québec, this fire season exhibited the driest conditions in 124 years and showed unprecedented drought severity across the province by mid-June (Vincent et al. 2020; ESCER 2023). The Eastern Subarctic and Eastern James Bay HFR zones experienced June DC values that surpassed or equaled historical records, indicating severe drought. Previous research (Girardin et al. 2009) highlighted a mid-20th-century decrease in extreme droughts in Canadian forests due to increased precipitation, but an increase in drought occurrences in northern forests due to warming. Since 2010, the Eastern Subarctic zone has seen more extreme drought events than in the 1970-1980s, aligning with earlier periods (1920-1960). The 2023 events may indicate an ongoing rise in drought severity in northern regions, with a pause in the 1970s. In the Eastern James Bay HFR zone, a post-1980s upward drought trend suggests a potential reversal of the declining trends observed up to 2002 (Girardin et al. 2009; Girardin and Wotton 2009). In this area, significant positive trends in summer DC values between 1950 and 2020 are observed, meaning an increase in drought conditions over southern shorelines of the Hudson Bay, from Manitoba towards Québec HFRs (see ESCER 2023). Similar trends are also observed in British Columbia (Parisien et al. 2023).

Over the last centuries, variations in drought severity and frequency have significantly influenced fire regimes in Québec. Research shows that fire activity is closely linked to temperature, precipitation, vegetation, and human activities, highlighting the complex relationship between climate change and fires (Carcaillet et al. 2001; Remy et al. 2017; Girardin et al. 2019). To understand the 2023 wildfires, it is essential to differentiate between the wildfire histories of the Intensive Protection Zone and Northern Protection Zone, as historical fire drivers differed between these zones.

In the Intensive Protection Zone, burn rates were high during the Little Ice Age (around 1250AD–1850AD; Gennaretti et al. 2014) and the early twentieth century (Drobyshev et al. 2017; Danneyrolles et al. 2021; Chavardès et al. 2022). However, they remained low from 1940 until the extreme 2023 season, coinciding with the growth of the modern forest industry and advancements in fire suppression (Boucher et al. 2017a; Tymstra et al. 2020). The 2023 fire season was the most active in commercial forests since 1923 (Figure 5a), but when compared to long-term records, it still falls within the natural range of variability (Chavardès et al. 2022). About 7.4% of the western part of the Intensive Protection Zone burned in 2023, contributing to a decade-long mean annual burn rate of ∼0.7%, which is still within the historical range of 0.6 to 1.3% from 1750 to 1950 (Chavardès et al. 2022).

Conversely, in the Northern Protection Zone, particularly its western portion, fire activity in 2023 differs significantly from its recent range of variability. This region is among the most pyrogenic in Québec and the circumpolar boreal zone. Employing a methodology by Héon et al. (2014), fire size was reconstructed using dendrochronology along transects (Figure 1). The fire regime within the Northern Protection Zone has been consistently active since 1800, marked by regularly occurring extreme fire years without a long-term trend (Figure 5b). Significant fire years in this region include 1847 (during which 90 km of the transects burned), 1882 (104 km), 1906 (104 km), 1922 (140 km), 1941 (96 km), 1989 (136 km), and 2013 (102 km). However, 2023 surpassed all these with 208 km burned transects, making it the most substantial fire year in the last 224 years.

Overall, these analyses confirm that the 2023 fire season is markedly distinct from those of the last century, in terms of severe weather conditions and areas burned, and that these conditions are getting more frequent in Québec. The fire activity, fueled by these conditions, is unprecedented in at least 220 years of record in the Northern Protection Zone. Despite being within the long-term natural range of variability within the commercial forests, the extent of these fires ranks among the highest of what has been recorded in the past century. Further global warming could exacerbate the ongoing trends identified above and ultimately lead to a 3- to 5-fold increase in annual area burned by the end of the 21st century in the province (Boulanger et al. 2014). Such increases in fire activity would strongly modify the boreal forest ecosystem (Chaste et al. 2019; Boulanger and Pascual 2021; Boulanger et al. 2022a), affect its ability to conduct sustainable forest management (Gauthier et al. 2015; Pau et al. 2023), substantially increase province’s fire management and suppression costs (Hope et al. 2016), increase infrastructure and communities’ exposure to short fire interval (Erni et al. 2021; Arseneault et al. 2023) and modify species-at-risk habitats (Tremblay et al. 2018; St-Laurent et al. 2022; Leblond et al. 2022). Impacts generated by the 2023 fire season and those projected in the upcoming decades because of climate change are presented in the following sections.

## 3. Impacts of the 2023 fire season

### 3.1. Effects on the forest sector

Québec’s forest sector, a key part of the province’s socio-ecosystem, faces significant challenges due to the 2023 wildfires, as well as the expected increase in such events in the coming decades. In 2020, this sector generated $CAN 20.6 billion in revenue and contributed $5.1 billion to the real GDP (NRCan 2020). Despite a decline over the past two decades (NRCan 2023), the sector remains crucial for the stability of many small, remote, and mono-industrial communities in Québec. The 2023 fire season induced important losses in silvicultural investments and will strongly impact timber supply over the coming decades. Drastic reductions in annual allowable cut (AAC) will occur in management units that were most hardly hit by the 2023 fires (Forestier en Chef 2023), which will span over several decades (see also Suppl. Mat. S3).

Furthermore, a conservative estimate of at least 300,000 ha of commercial forests might suffer from regeneration failures (Le Devoir 2023a; see Suppl. Mat. S4) because these stands were immature (<60 years) with an insufficient regeneration potential (Splawinski et al. 2019). Unless these areas are planted, they will likely remain unproductive for decades. These young stands are primarily associated with the cumulative historical impacts of harvesting and wildfires over the last 50 years (see Suppl. Mat. S4 - Table S4.2). Considering the current capacity of the forest sector for plantation (∼50 kha per year), bringing back these areas into production could take several years, a huge budget (probably several billions of dollars) and imply technical challenges at a level never seen before (e.g., building of new forest roads, shortages in seedlings, nursery, and planting labor). Economic consequences of these impacts reside in the substantial losses in silvicultural investments, including many plantations. We estimate that ∼80,000 ha of plantations have burned in 2023 (Table S4.2).

To alleviate fire-induced losses, burnt forests with mature trees can be targeted for salvage logging. However, considering the extent of 2023 fires, and that mature forests represent a fraction of what has burned, only ∼10-20% of burned stands are likely to be salvaged in the upcoming year. As such, salvage logging will not offset the long-term deficit in timber supply induced by the 2023 fires. Additionally, recently burned trees sought by salvage logging are rapidly affected by degrading agents (e.g., wood boring insects and checking), that reduce the economic value of postfire timber, and thus the profitability of coping practice (Saint-Germain and Greene, 2009; Boucher et al. 2020). Salvage logging, following a wildfire, acts as a second disturbance in a brief period. This combination of fire and logging can significantly alter the recovery and provision of various ecosystem services (Lindenmayer and Noss 2006; Leverkus et al. 2020). Salvage logging affects numerous aspects like tree seedbeds, seedling density, woody debris, biological legacies, water conditions, and soil properties (Leverkus et al. 2018, and 2020). These influences have short- to long-term effects on the forest’s structure, composition, diversity, and dynamics (Purdon et al. 2004; Nappi et al. 2011; Thorn et al. 2018).

The immediate and extended impacts of the 2023 fires on Québec’s forest sector reflect the numerous warnings issued by the scientific community in the past two decades. It was shown that climate-induced increases in wildfire activity posed a serious threat to achieving sustainable forest management goals (Gauthier et al. 2015; Forestier en Chef 2021), with wood shortages and regeneration failures representing a primary climate-induced risk for Québec’s forests. With the expected surges in wildfires, Québec’s commercial forests will face an increasing proportion of immature forest at the expense of harvestable forest stands, despite a possible increase in productivity in the northern forest (Pau et al. 2022; Wang et al. 2022; Danneyrolles et al. 2023). In addition, simulation experiments suggest that enhancing vegetation productivity may result in a positive feedback loop with increased fire activity, ultimately yielding little net benefit over the long term (Chaste et al. 2019). Business-as-usual strategy could imply permanently lowered timber supply (by more than 60% in some regions) to avoid fire-induced timber shortages (Forestier en Chef 2021). An increasing fire activity is also likely to amplify broadleaved tree regrowth at the expense of the preferred conifers (Boulanger and Pascual 2021), increasing salvage logging (Forestier en Chef 2021) and management costs (Cyr et al. 2022). This scenario could lead to a “manager’s dilemma,” where the significant social costs of intensively managing forests, such as replanting conifers in areas with failed post-fire regeneration, must be balanced against the increased vulnerability of these forests to future wildfires. Also, frequent softwood timber supply shortages, higher stumpage prices and decreased export value of forest-related products could seriously affect the province’s economy and potentially prompt the devitalization of forest communities in the long term (Williamson et al. 2007).

### 3.2. Impacts on the fire management agency, communities, and infrastructure

With an overwhelming number of simultaneous and intense wildfires, the 2023 fire season really put at test Québec’s fire management agency’s (SOPFEU) operational limits, which are known to cap out at about 30-40 active fires a day. Indeed, in the Intensive Protection Zone only (Figure 1; also refer to Cardil et al. [2019]), the number of active fires jumped from 21 to 132 on June 1st, marking the start of 45 consecutive days with over 30 active fires. These numbers greatly exceed the historical (1994-2022) average of 14 ± 17 (mean ± SD) days with over 30 active fires per year. As a result, the exceeding fires burned freely until reinforcements from the Canadian Armed Forces, and other provincial and international fire management agencies started to arrive on June 5th, adding 50 people to the 434 local resources already at work. External reinforcements peaked on June 28th with 994 people, while local resources peaked at 643 on June 12th. The lack of readily available resources to respond to the workload of early June, coupled with the inefficient or unsafe conditions of fire suppression activities due to the extreme fire weather (Hirsch and Martell, 1996), likely contributed to rapid spread of these fires, sometimes threatening communities and infrastructures. This led SOPFEU to prioritize fires threatening human lives and/or infrastructures deemed essential to public security, that were numerous, over those simply threatening the forest, including silvicultural investments and standing timber (Cardil et al. 2018).

As climate change intensifies, scenarios like the 2023 fire season, where fire management capacities are overwhelmed, are expected to become more frequent. This is due to projected increases in fire occurrences, area burned, conditions favorable to fire spread, and days with intense fires that impede suppression efforts (Wotton et al. 2017). Recent work indicated that this would have a direct impact on both wildland firefighters and airtankers workload (i.e., number of hours worked doing fire suppression activities) (Boulanger et al. 2022b; Boucher et al, in preparation). The 2023 fire season is an example of these impacts, with a total firefighters (local and external) workload that summed up to 755,648 hours, representing more than six times the historical (1994-2018) average of 117,017 ± 121,530 (mean ± SD) hours, and more than 1.5 times the maximum of 477,024 hours observed in 2010 (Boucher at al, in preparation; 2023 data provided by SOPFEU). Airtankers flew a total of 3,219 hours fighting fires in 2023, more than three times the historical (1994-2018) average of 981 ± 740 (mean ± SD) hours, and just over the observed maximum of 2005 with 3,096 hours (Boucher at al, in preparation; 2023 data provided by SOPFEU). The 2023 fire season observed workload for both firefighters and airtankers fall within the range of expected future workload for the end of this century (2071-2100) under RCP 8.5, under which 474,300 ± 211,680 (mean ± SD) hours for firefighters and 5,832 ± 4,212 (mean ± SD) hours for airtankers are projected (Boucher et al. in preparation). As extreme fire-prone conditions are expected to become more frequent, the 2023 wildfire season is thus a reminder that costs associated with fire management and suppression will greatly increase in the upcoming decades (Hope et al. 2016), thus putting more communities and infrastructures at risk.

Many communities faced direct fire threats and had to swiftly react to fires that were rapidly progressing towards critical infrastructures during the 2023 fire season. As a last barrier of defense, approximately 45-69 km (or 226 ha) of fire breaks (where vegetation was stripped to the mineral soil on a width of approximately 50 m) were created in a hurry to directly protect seven communities (Chapais, St-Lambert, Normétal, Chibougamau, Oujé-Bougoumou, Mistissini and Lebel-sur-Quévillon, data provided by the *Direction de la Protection des Forêts* of the *Ministère des Ressources Naturelles et des Forêt du Québec*). These conditions also led to an unprecedented number of wildfire-related evacuations, with over 38,700 people evacuated, including many from First Nation communities (CFS 2023). Large northern Québec communities like Sept-Îles (population of about 25,000) and Chibougamau-Mistissini (population of about 10,000) were among those evacuated. Some towns, such as Lebel-sur-Quévillon, experienced multiple evacuations, adding stress to the residents, local public safety agencies, and officials. On June 9th, the province announced a financial help of 1,500$ to each evacuated household (Gouvernement du Québec 2023a). Based on the number of evacuees per communities and the number of people per household (Statistics Canada 2023a) we estimated that about 14,509 residences were evacuated, for a total toll of this financial help estimated to 21.8 M$. The mental health toll was evident, as seen in the resignation of the mayor of Chapais in November 2023 due to post-traumatic stress disorder caused by the intense wildfire crisis management (La Presse 2023). Remarkably, there were no casualties directly linked to the fires. However, there were significant losses in terms of infrastructure, including forest machinery (Le Soleil 2023) and First Nation’ infrastructures on traditional territories (Grant 2023). Indeed, approximately 1,154 structures (min = 1, max = 627 structures per fire; MCBF [2020] and Gouvernement du Québec [2023b]), mainly cabins, were in the path of 64 wildfires this season, leading to the destruction of many (J. Boucher, personal observations).

The significant wildfire activity in the James Bay region (Figure 4b) consistently poses threats to communities and crucial infrastructures in Northern Québec, particularly the La Grande hydroelectric complex, a vital part of the province’s energy network (Erni et al. 2017; Arseneault et al. 2023). This complex, contributing 40% of Québec’s electrical power, comprises eleven hydroelectric stations and related infrastructures like high-voltage power lines, roads, airports, and residences. In 2023, many fires impacted strategic areas of Hydro-Québec, crossing major power lines that serve the province’s most populated regions (Figure 1), resulting in multiple shutdowns of high-voltage lines (Le Devoir 2023b). A similar situation occurred in 2013, leading to power outages affecting major areas, including Montreal, Québec’s largest city, disrupting the subway system and affecting hundreds of thousands of people. With anticipated increases in fire frequency and intensity due to climate change, concomitant with growing population and energy demands, hydroelectric installations and transportation networks in this region face growing risks (Arseneault et al. 2023), emphasizing the need for enhanced fire risk mitigation near these critical infrastructures in remote areas.

**Figure 4.**
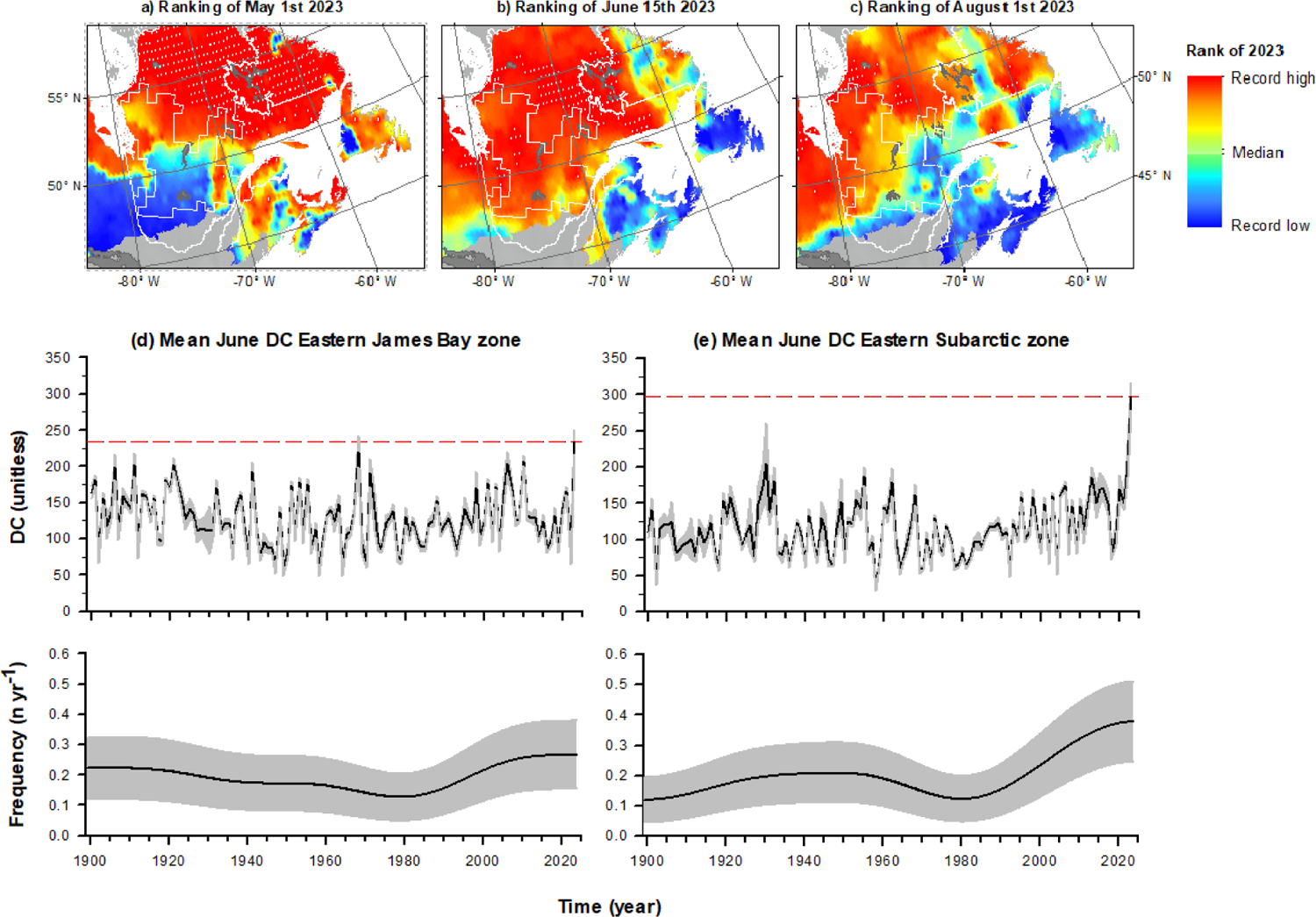
Drought Code (DC) severity trends in Québec’s boreal forests for Early-May, Mid-June, and Early August (1900-2023). Map resolution: 0.5 degrees. (a) Ranking of DC severity for May 1st, 2023, against the historical DC severity for the same date over a 124-year period. Regions in red represent instances where the daily DC level in 2023 closely approached the historical severity level at each grid point (refer to the legend on the right), while white dots denote record-high levels for 2023. (b-c) Similar to (a) but focusing on June 15th and August 1st. (d-e) Middle-row plots: Time series illustrating the daily DC average for June encompassing the Eastern James Bay and Eastern Subarctic zones. The gray shading represents 90% confidence bands accounting for spatial autocorrelation. Horizontal dashed lines mark the severity of 2023. Bottom-row plots: Analyses of extreme DC severity trends, estimated as the rate of occurrence (per year) of extreme drought years using a kernel approach (bandwidth parameter h=15; based on Mudelsee 2002; Mudelsee et al. 2004). The gray shading represents 90% confidence bands for risk estimates. Additional details regarding the methodology are available in the Supplementary Information.

**Figure 5.**
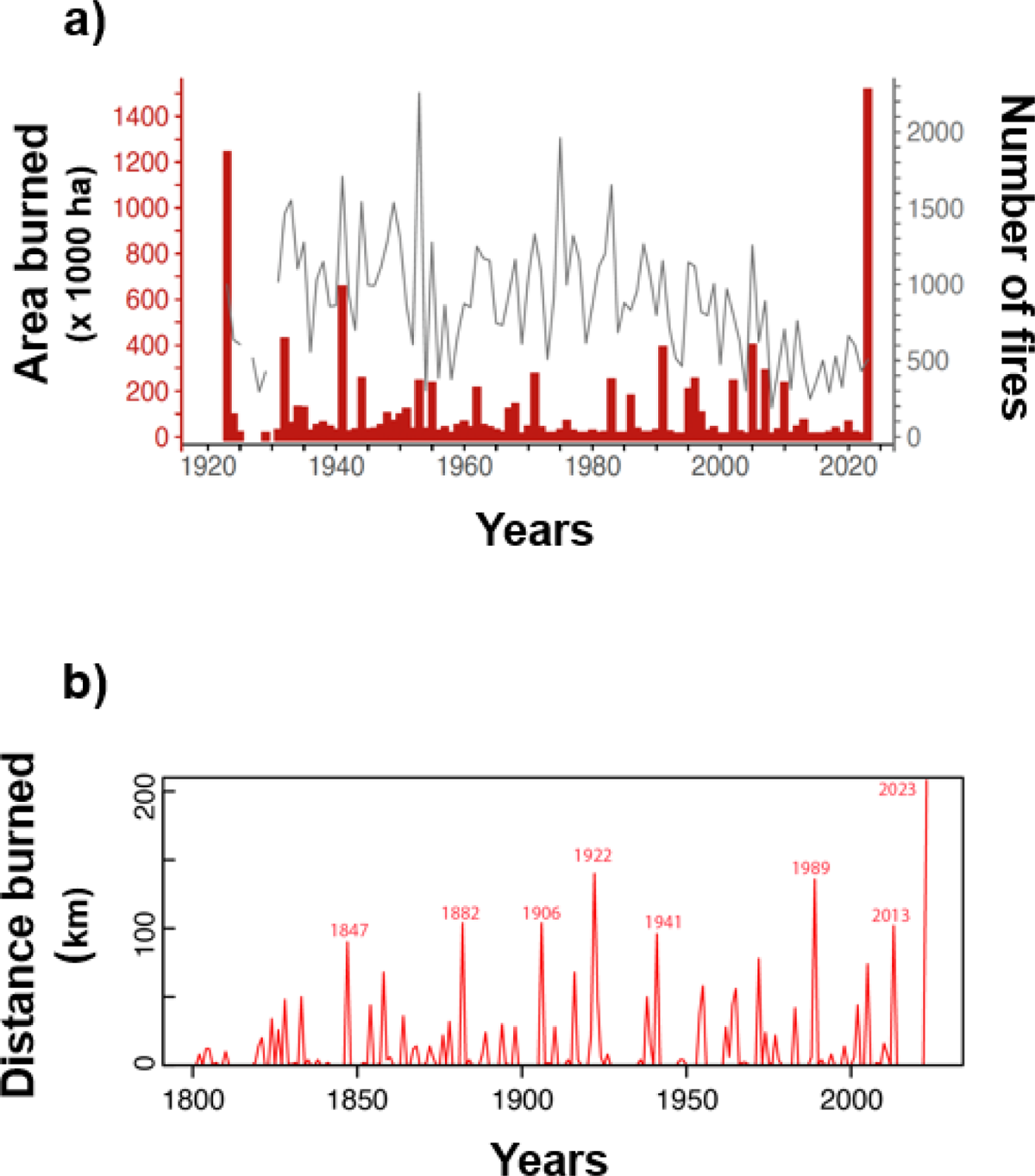
**a)** Annual area burned (AAB) and number of fires within the Intensive Protection Zone (IPZ) in Québec between 1923 and 2023 (retrieved from the archives of the *Ministère des Ressources Naturelles et des Forêts du Québec*); b) Distance burned along the Billy Diamond and Trans-Taiga roads in northern Québec (total of 640 km sampled). The years indicated correspond to fire events covering 90 km or more.

### 3.3. Impacts on wildlife and their habitat: the boreal caribou as a case study

The 2023 Québec wildfires are expected to impact wildlife differently depending on species: burn-associated species (Boucher et al. 2012; and 2016) or species adapted to early seral or open forests may benefit (Hutto and Patterson 2016; Knaggs et al. 2020), while those reliant on old-growth forests could suffer due to habitat loss exacerbated by industrial forestry and climate change (Drapeau et al. 2016; Bergeron et al. 2017; Rudolph et al. 2017; Tremblay et al. 2018). In Québec, boreal populations of woodland caribou (*Rangifer tarandus*; hereafter caribou) are respectively listed as threatened and vulnerable under the federal and provincial species at risk acts. Cutovers originating from industrial timber harvesting and associated road networks are regarded as the main causes of caribou population declines in the province (MFFP 2021).

Environment Canada (2011) showed that caribou demography was best explained by a combination of wildfire and anthropogenic disturbances, with populations living in heavily disturbed ranges having a lower recruitment rate, and to a lesser extent, a lower adult survival (Johnson et al. 2020). Considering the uniqueness of the 2023 fires, we sought to determine how much of the caribou’s distribution range had burned, and we explored the contribution of recent fires on the area covered by total disturbances for several caribou ranges.

Across the 11 caribou ranges studied (Fig. 6), the 2023 wildfires burned 1,490,100 ha, representing on average 2.6% ± 3.7% (SD) of range areas (Fig. 7a). The three ranges occurring in northwestern Québec had the highest proportion of area burned, with the Nottaway range showing the highest proportion (11.3%). Before 2023, caribou ranges had on average 43.9% (± 27.1%) of their area covered by total disturbances, defined as the sum of natural disturbances (0-40 years old burned areas) and non-overlapping anthropogenic disturbances (0-50 year-old clearcuts and roads buffered by 500 m). Most human disturbances occurred south of the northern forest allocation limit, i.e., within the commercial forests. After the 2023 wildfires, caribou ranges were disturbed at 48.0% (± 26.8%) on average, representing a 14.0% (± 22.7) increase (i.e., rate of change) compared to pre-2023 conditions (Fig. 7b). Some fires occurred in young regenerating forests, which were already considered as disturbed caribou habitat. These fires had the consequence of burning the regeneration, bringing the areas back to a younger disturbed state, but they did not add to the percent total disturbances in our analyses.

**Figure 6.**
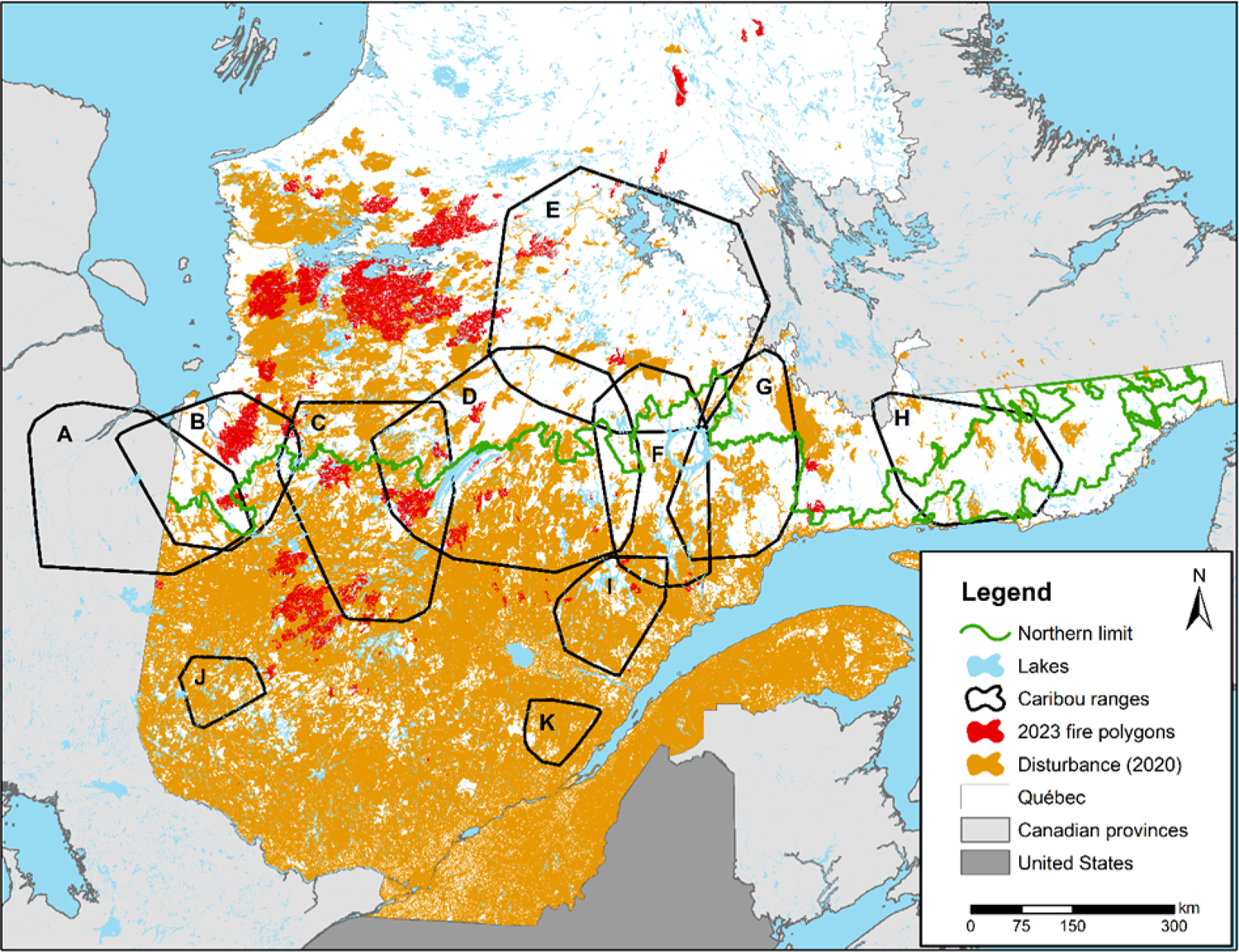
Location of the 11 boreal caribou ranges (black polygons) (A: Detour; B: Nottaway; C: Assinica; D: Témiscamie; E: Caniapiscau; F: Outardes; G: Manicouagan; H: Basse-Côte-Nord; I: Pipmuacan; J: Val d’Or; K: Charlevoix). The map also shows, in orange: the area covered by total disturbances (natural and anthropogenic) as identified in 2020 using the most recent 1:20,000 ecoforest maps published by the Québec government; in red: the 2023 wildfires. Natural disturbances include fires (0-40 years old). Anthropogenic disturbances include roads and clearcuts (0-50 years old), buffered by 500m. The northern forest allocation limit above which commercial timber harvesting is prohibited is shown using a green line.

**Figure 7.**
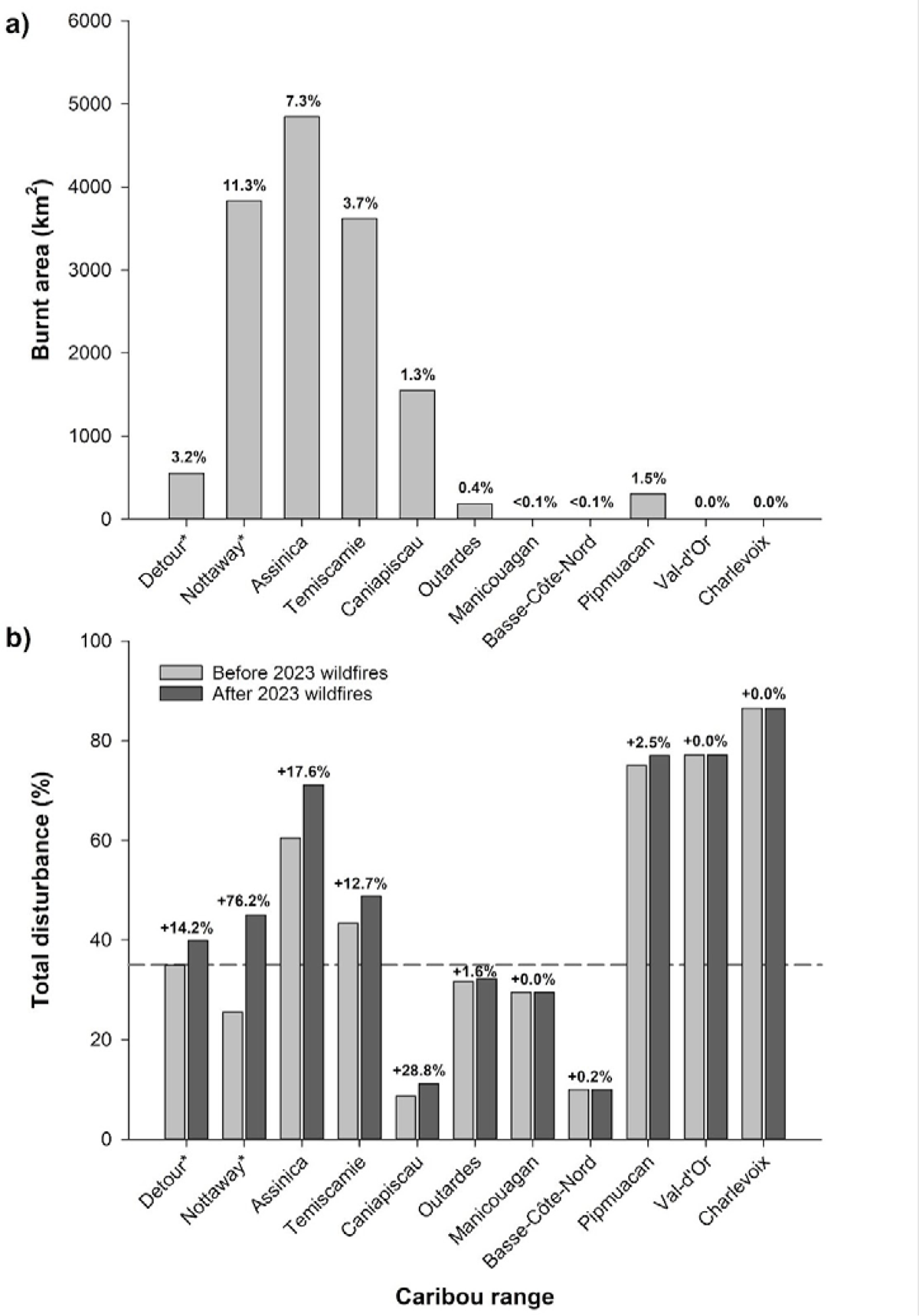
**a)** Area (in km^2^, gray bars) and percentage (numbers above bars) of boreal caribou ranges that were burned by the 2023 wildfires. **b)** Histogram synthesizing the total disturbance levels in boreal caribou ranges before (light gray) and after (dark gray) the 2023 wildfires. The dashed line represents the 35% disturbance management threshold used by Environment and Climate Change Canada to identify populations more likely than not to be self-sustaining in the long term. Percentages above the histogram bars represent the rate of change in total disturbances caused by the 2023 wildfires (i.e., the newly impacted areas), compared to the pre-2023 disturbance levels. *These ranges extend to Ontario; only the Québec portion was analyzed.

Wildfires in 2023 increased total disturbances in 8 of 11 caribou ranges in Québec. These fires brought the Detour and Nottaway ranges above the 35% management threshold set by Environment Canada (2012), i.e., the threshold above which a population has less than a 60% probability of being self-sustaining in the long term (Fig. 7b). Before the 2023 fires, 5 of the 11 ranges were already above this threshold. Studies by Environment Canada (2011) and Johnson et al. (2020) have indicated that wildfire impacts on caribou were less severe than those of human disturbances. Caribou are adapted to the dynamic boreal ecosystem, regularly affected by forest fires of varying frequencies and severities (Lafontaine et al. 2019). Therefore, the 2023 fires’ additional impacts on caribou were likely modest province-wide, although some populations were more affected than others. Should salvage logging operations be initiated within the distribution range of caribou, the consequences could be detrimental for the species, notably if new permanent roads are introduced into the landscape (Lindenmayer and Noss 2006).

Additionally, the cutblocks that would be created following salvage logging would exert a more severe negative impact on caribou populations compared to burned areas (Johnson et al. 2020). In this context, on 8 November 2023 the Québec’s Ministry of Natural Resources and Forests prohibited salvage logging operations within a 16,700-ha fire located within the Pipmuacan caribou range overlapping a projected protected area under the stewardship of the Pessamit Innu First Nation.

A sound management and conservation strategy for caribou is urgently needed in Québec. This strategy should implicitly consider the impacts of fire seasons such as that of 2023, expected to occur more frequently in the future (Boulanger et al. 2014). It should also tackle the main cause of caribou declines, i.e., landscape disturbances originating from industrial timber harvesting (St-Laurent et al. 2022; Morineau et al. 2023). Conservation approaches focusing on habitat protection (Leblond et al. 2022) and restoration (Lacerte et al. 2021, 2022) should benefit caribou, as well as other species of the boreal forest (Bichet et al. 2016), although complementary conservation actions would be needed to cover a broader extent of species (Micheletti et al. 2023).

### 3.4. Impacts on First Nation communities, territories, and people

First Nation communities have historically been facing disproportionate consequences of wildfire: many are located within the wildland-urban interface, are exposed to greater fire risks, and are overrepresented in wildfire-related evacuation events (Erni et al. 2021). This trend actualized in 2023, with Eeyou (Cree), Anishnaabe (Algonquin), Atikamekw and Innu communities experiencing the consequences of wildfires. According to primary data collected by W8banaki emergency center, who have supported First Nations during the 2023 fire season, more than 10,000 people within thirteen communities were evacuated. Threats were both direct and indirect and included risks for people and infrastructures, bad air quality over extended periods of time, and closures of roads that were often the only way of accessing the communities. The communities’ vulnerability exposed in 2023 is consistent with previous assessments showing that the transition of the Eeyouch from a traditional lifestyle based on trapping, hunting, fishing, and gathering to a more sedentary way of life has made their new infrastructures more vulnerable to fire (Arseneault et al. 2023).

The wildfires of 2023 have also dramatically transformed First Nation territories. The Eeyou and Anishnaabe family hunting grounds, located in the western boreal zone, were especially affected. From the 384 Eeyou, Abitibiwinni and Lac Simon hunting grounds, 119 (31%) had more than 5% of their land surface burned in 2023. Thirty-one hunting grounds had more than 50% and nine had more than 80%. Hunting grounds are land units passed down from one generation to the next that are key places for cultural and subsistence practices including trapping, hunting, fishing, speaking language, learning bush skills, and celebrating important life events. The losses experienced by numerous families were thus tremendous, both economically and culturally.

The effects of the 2023 wildfires on First Nation territories accumulate with previous (and future) disturbances from industrial activities, mostly through forestry, mining, and hydroelectric development (Bélisle and Asselin 2021). For instance, forty-three hunting grounds were affected both by 2023 wildfires and by timber harvesting (1990-2020) and are likely to be subjected to salvage logging. As salvage logging operations stem from special plans, they often involve derogations that allow operations to deviate from existing regulations regarding forest roads or the protection of regeneration, for instance, and do not need to undergo the regular public consultation process (Gouvernement du Québec 2023). Consequently, consultations with First Nations and other regional and local stakeholders are rushed, with only a few weeks to assess and anticipate the impacts of thousands of hectares of salvage logging.

First Nation people have been coping and adapting to wildfire for centuries, developing fine knowledge of burned forests and their resources (Miller et al. 2010). However, for highly disturbed hunting grounds, adding wildfire to previous changes initiated by industrial activities may surpass people’s adaptation capacity and affect their quality of life and general wellbeing (Parlee et al. 2012; Fuentes et al. 2020). Even for hunting grounds with low previous disturbance levels, salvage logging could cause rapid and cascading changes, including increased traffic, industrial development, and land use conflicts (Walker et al. 2011; Bélisle and Asselin 2021).

Moreover, the rigid land boundaries between family hunting grounds that were established within the James Bay and Northern Québec Agreement has reduced families’ adaptability to large scale disturbances by limiting their access to the entire land (Sénécal and Égré 1999). The interplay between industrial activities and climate change is significantly reshaping the environmental and cultural landscapes in which the First Nation people live, and the extreme fires of 2023 are set to become a defining part of this transformation.

## 4. Where do we go from here?

The Québec forest industry has faced several crises over the years, including the severe spruce budworm outbreak of the 1970s and 1980s, the softwood lumber crisis (2005-2010), and the *Commission d’étude sur la gestion de la forêt publique Québécoise* (2004). Each time, the industry had to adjust its practices and adapt to the evolving situation. Likewise, Québec has faced important climate-related catastrophes in the last decades (e.g., the 1998 ice storm, the major floods of 1996, 2011, 2017, 2019, and 2020; see the list of all natural disasters in the Canadian Disaster Database; Public Safety of Canada 2023), that mandated profound changes to the way society operates (for example, see the “*Plan de Protection du Territoire Face aux Inondations*” developed by the Québec government in April 2020 in response to the 2019 flood event, [Gouvernement du Québec 2020]). The 2023 wildfires are among these major crises that will pose a substantial challenge for the upcoming years, highlighting the urgency to adapt and reduce the risks associated with future fires. We propose some solutions below.

### Solution 1: Considering fire *a priori* in the calculation of the annual allowable cut

When considering overall impacts on timber supply and biodiversity, current forest management in Québec is maladapted to face climate change. Harvesting rates may be just too high to sustain a steady timber supply, especially under the new fire regime. One of the most effective and immediate large-scale adaptation strategies would be to create precautionary wood reserves. As a corollary, this implies factoring-in wildfire impacts upfront when calculating the annual allowable cut (AAC). In Québec, fire effects are typically factored into the AAC *a posteriori* (with very few exceptions). This means that after a fire occurs, the AAC of the affected forest management unit is recalculated to account for the long-term changes in the availability of harvestable stands caused by fires. In accordance with this strategy, Québec’s Forester in Chief has recommended in November 2023, a 12.7%, 2.1% and 0.2% decrease in AAC within the three regions most affected by the 2023 wildfires (Nord-du-Québec, Abitibi-Témiscamingue and Mauricie) (Forestier en Chef 2023). However, many studies have shown that such a strategy could result in important variations in AAC over time and would impede sustainability in the long term (Raulier et al. 2014; Leduc et al. 2015; Daniel et al. 2017; Forestier en Chef 2021). This prevents the industry from benefiting from predictability in the AAC to plan future activities. Depending on the current and future fire regimes, an *a priori* consideration of wildfires could lead to the establishment of precautionary wood reserves representing e.g., ∼5 to 20% of the AAC. Although this could initially be seen by some as a negative measure, it would in fact increase the probability of maintaining a constant long-term timber supply predictability, which would be beneficial for the industry in general (Savage et al. 2010). Considering precautionary reserves upfront in the calculation of the AAC would help prevent shortfalls and *a posteriori* reduction in timber supply for regional burning rates as low as 0.30% to 0.45% (Savage et al. 2010; Ministère des Ressources Naturelles du Québec 2013). A preliminary analysis conducted for the current study (Suppl. Mat. S3) shows that, if a 20% precautionary reserve had been established 20 years ago in northwestern Québec, the 2023 wildfires would have not led to any drastic postfire decreases in timber supply. Conversely, such a precautionary reserve would have led to more timber harvested in the medium to long term (>2030-2040). Burning rates that would require the establishment of precautionary reserves are already affecting vast areas of the commercial boreal forest in Québec. By mid-century, even under moderate climate change, these rates are expected to impact nearly all the managed boreal forests in Québec (Boulanger et al. 2014; Pau et al. 2023), adding to the urgency of establishing these reserves. Maintaining precautionary wood reserves would also benefit several additional ecosystem services, notably by maintaining old-growth forests that are high-quality habitat for many wildlife species including caribou (Bichet el. 2016; Leblond et al. 2022; St-Laurent et al. 2022; Labadie et al. 2023).

### Solution 2: Making forest landscapes more resilient to fire

Increasing forest resilience to wildfires could help reduce postfire regeneration failures. This could be achieved notably by favoring species with an early sexual maturity such as jack pine (Rudolph and Laidly 1990; Cyr et al. 2022). Variable retention or partial harvesting leaving mature trees after logging could also be envisioned in fire prone black spruce-jack pine-dominated landscapes: spared trees in a dispersed or aggregated pattern could serve as seeding trees if a burn was to occur a few decades after logging (Perrault-Hébert et al. 2017; Cyr et al. 2022). Perrault-Hébert et al. (2017) showed that leaving between 10 and 15% of mature seed trees could be sufficient to restore a low to moderate level of regeneration and avoid the high social costs of post-fire plantations (5-8k$.ha^-1^, based on 2023 estimations). Seed tree retention could also mitigate the impacts of severe mature biomass removal on forest biodiversity (Thorn et al. 2020). The precautionary wood reserves discussed above would also mitigate the increase in regeneration failures by leaving more mature stands at the landscape scale. As fires are promoting hardwood species, there’s also a need to reevaluate post-fire forest management strategies that favor coniferous species over time. Increasing functional redundancy, through specific forest management and silvicultural practices, could also help increase forest resilience after disturbance and foster the provision of ecosystem services (Messier et al. 2019).

### Solution 3: Making forest landscapes more resistant to fire

An additional strategy would aim at making forest landscapes more resistant to climate-induced increases in wildfire activity and to reduce their consequences. Higher resistance could stem from changing, either actively or passively, the flammability of the vegetation, notably through increasing the hardwood component of forest landscapes (Terrier et al. 2013). When fully leafed, hardwood species such as aspen, white birch, and red maple are known to be less flammable than conifer species such as balsam fir, black spruce and jack pine (Forestry Canada Fire Danger Group 1992; Hély et al. 2001, 2010; Bernier et al. 2016). Simulations have shown that actively planting or favoring the natural regeneration of hardwood species after fire or harvest could strongly alleviate the climate-induced increases in fire activity and mitigate concomitant losses in timber supply by 50% (Forestier en Chef 2021). Preliminary analyses (Suppl. Mat. S4) revealed that hardwood stands, more than any other types of forest cover, were underrepresented in the forested areas burned in 2023, underscoring the potential protective qualities of hardwood stands even during severe wildfire seasons.

However, there are several drawbacks to this strategy. The protection ability of hardwood trees is limited before leafout, a period during which a significant number of fires can occur within the boreal forest (Parisien et al. 2023). Furthermore, not all boreal sites can support hardwood species due to specific soil characteristics (Marchais et al. 2022). In the western boreal bioclimatic domain of Québec, hardwood and mixed stands occupy just under 10% of the forest area, primarily on hillsides (Blouin and Berger 2004). The protection ability of hardwood should thus be assessed in the light of the ecological classification of Québec’s forest ecosystems (Saucier et al. 2010). In addition, the operational capacity of the forest sector to convert forest landscapes is limited in space and time. In this context, it might be advantageous to consider how the 2023 fires will themselves alter forest landscapes by increasing the pioneer hardwood components in the forests (Boucher et al. 2014, 2017b).

In addition to these drawbacks, a rapid anthropogenic conversion of boreal landscapes would have tremendously deleterious impacts on a myriad of species associated with conifer forest covers (Tremblay et al. 2018; Labadie et al. 2023), species that are also typically vulnerable to logging (Imbeau et al. 2001; Venier et al. 2014; Leblond et al. 2022) and climate change (Bouderbala et al. 2023). Rapid conversion of forest landscapes could also significantly alter the livelihoods, cultures, and identities of First Nations who are closely tied to the land (Belisle et al. 2022). In this context, such a strategy could be limited to local conversion of forest landscapes by aiming to decrease wildfire risks to communities or critical infrastructures.

Alternatively, the valuation of wetlands as fire breaks and biological refuges through conservation has been exposed as a contributor to forest resistance and resilience to wildfire in western United States (Fairfax and Whittle 2020). Wetlands provide several ecosystem services (Cimon-Morin et al. 2016) and are of primary importance to culture and subsistence for First Nations, particularly for hunting (Grant 2024), as well as being a passive and inexpensive method of increasing the fire resistance of forests. This unexplored solution could be an opportunity for further collaborative research on wildfires on First Nation lands. However, it is important to note that vegetated wetlands may also carry fire during drought conditions (Canadian Forest Service Fire Danger Group 2021), which are expected to become more prevalent with climate change.

### Solution 4: Adapting the forest management system and the industrial structure

The 2023 fire season raises questions about the forest sector’s ability to adapt to extreme fire events (Boulanger et al. 2023). Forestry practices, which are based on ecological classifications such as potential vegetations, will certainly have to be revised as climate change itself will impact postfire successional pathways at each ecological classification level (Grondin et al. 2022). On the operational side, the ability of the forest sector to intervene on the landscape is limited. For instance, it might be difficult to restore all the current and future postfire regeneration failures, due to limitations in budget, labor or nursery capacity. Moreover, the capacity of the forest sector to salvage wood, despite its special plans, will remain logistically limited, and a significant proportion of the burned stands will be left without active management. This underscores the importance of proactively preparing the forest to withstand more frequent fires. The large area burned in 2023 might also represent the opportunity to test alternative management strategies to increase the resistance and resilience of forest landscapes to wildfires and climate change impacts.

Adapting the industrial structure will be paramount to make the whole forest sector more resilient. For example, a significant increase in hardwood content (i.e. aspen and white birch) within the timber supply following increased fire activity might prompt a significant paradigm shift in a forest sector that traditionally relies mostly on conifer species (Brecka et al. 2020). Increased fire activity could lead to novel uses of salvaged wood, such as pulpwood or bioenergy. Adapting the industrial structure might be more efficient than reactively adapting the forest to rising fire activity, although this will imply developing skills and capacity for action within communities. Innovation in this regard will be crucial, and incentives to promote it will have to be prioritized. In any case, it is likely that the industrial sector will need to focus on anticipating these changes, rather than merely reacting to them, to enhance its own vitality as well as that of the forest communities.

### Solution 5: Increasing suppression capacity and mitigating risks to communities and infrastructures

Even after considering all of the solutions proposed here, the fire management agency’s (SOPFEU) operational capacity of 30-40 active fires per day will likely have to be increased if we are to reduce the number of fires that are freely burning each year. On 14 November 2023, the government of Québec announced a 16 M$ investment in SOPFEU, for 2023-2024, to support increased suppression and prevention capacity (SOPFEU 2023b). This is a step in the right direction, but it may not suffice. Aerial suppression capacity seems to have capped out at just over 3,000 hours per fire season, as proven by the fire seasons of 2005 and 2023. This could be due to the worldwide aging fleet (Radio-Canada 2023) of the airtankers that are most efficient in our boreal conditions (mainly CL-215 and CL-415, McFayden et al 2023) and the scarcity of qualified pilots (Noovo Info 2023). New aircrafts would be welcome, and governments may need to commit to purchasing more aircrafts to ensure that they are ready for service by 2030 (Le Soleil 2023). In parallel, new pilots and mechanics will also need to be recruited and trained.

This fire season, lots of resources (staff and machinery) that could have otherwise been tasked to suppression operations were requested to support the fast and reactive responses to safeguard communities and infrastructures from the wildfires, highlighting the essential need for more proactive risk management near sensitive areas. This includes considering risk assessments to identify areas likely to burn and implement mitigation measures around corresponding communities and infrastructures before an emergency occurs. It would also enhance the level of awareness and preparedness among communities in addressing wildfire-related emergencies. For example, antecedent mapping of burn probabilities in the La Grande Rivière hydroelectric complex in the James Bay area could identify the areas that preferentially burned in 2023 (Arseneault et al. 2023), demonstrating the accuracy and usefulness of such predictive assessments in wildfire management (Figure 8; Parisien et al 2019). Maintaining an organized database that records the impacts of wildfires on infrastructures is also vital. Such a database would provide valuable insights into the extent and severity of damage and the effectiveness of existing mitigation strategies. This could inform future risk analyses by providing data to develop susceptibility functions for resources and assets threatened by fire, adding to those developed for residential structures (Abo El Ezz et al. 2022; Nicoletta et al. 2023). Furthermore, it is important to increase the monitoring capacity, encompassing aspects like evacuation procedures and the resistance of infrastructures. Enhanced monitoring (e.g., through daily remote sensing) would facilitate timely responses during wildfire emergencies, potentially saving lives and reducing property damage. The adoption of FireSmart practices (CIFFC 2023) is a key part of this strategy. In this regard, communities that have established fire breaks in a hurry this year, are now faced with crucial decisions regarding the future of these protective measures. A potential solution could involve intensifying land management near communities, for example by planting less flammable, fast-growing tree species such as hybrid poplar and hybrid larch, or shrub species like willow and alder, which could also serve purposes like biofuel production (Mansuy et al. 2018). Such measures not only safeguard against the immediate threats of wildfires but also contribute to the long-term resilience and sustainability of these regions. Cognizant of the need for more organized wildland fire risk management, SOPFEU, right after the fire season 2022, created a risk mitigation branch, while the *Ministère de la Sécurité publique du Québec* recently announced 31 M$ to implement mitigation measures such as fuel treatment and FireSmart measures in communities at high risk of fires.

**Figure 8.**
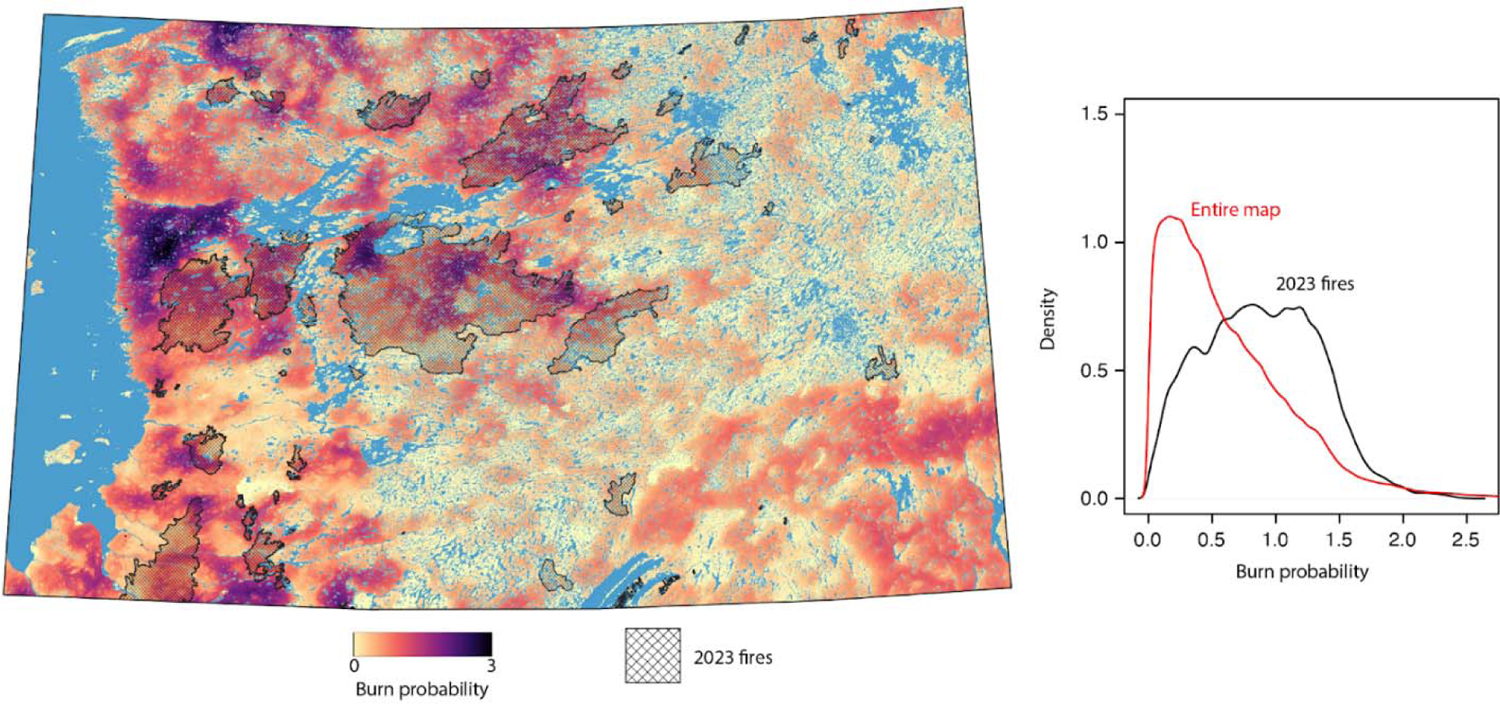
Comparison of areas burned in 2023 with fire probabilities mapped in 2022 across the strategic La Grande hydroelectric complex in the James Bay area. Map adapted from Arseneault et al. 2023.

First Nation communities had to mobilize very quickly to respond to the forest fires of 2023. They updated their emergency response protocols and developed their knowledge of fire risk assessment and fire behavior. They gathered information daily to make decisions with far-reaching consequences for the health and safety of their members. The people who were at the heart of the crisis in the communities were key players in the development of risk assessment and management tools for future fires. In this way, the 2023 fire season may have generated an opportunity for collaboration between scientific and First Nation institutions, both of which possess complementary knowledge and skills.

### Solution 6: A call for a unified risk management approach

The lessons learned from the 2023 forest fires in Québec also constitute an opportunity to review or revisit risk management strategies, in the context of on-going climate change and the occurrence of systemic risks and of tipping points (see the Global Assessment Report, UNDRR 2022). First, it is important to recall that a risk is not exclusively due to hazards, i.e., forest fires and their changes under warming conditions, but include other components such as human and natural factors. In fact, climate risks are the result of dynamic interactions between meteorological hazards and climate extremes, pre-existing local (environmental and human) vulnerability and exposure factors, as well as the collective and individual ability to cope (see IPCC 2018, 2023; UNDRR 2019 and 2022). Exposure and contexts of vulnerability and resilience affect the consequences of climate change and its associated risks (e.g., Berry et al. 2010; IPCC 2012; Disse et al. 2020). Both exposure and vulnerability factors change rapidly with socioeconomic and demographic developments, whereas forest fire hazards, in terms of both occurrence, duration and severity/extent will increase over the following years (Boulanger et al. 2023; Barnes et al. 2023). Hence risk management needs to evolve rapidly and continuously. This is one of the most challenging considerations for decision makers, practitioners, and industrial forestry managers, and this also means that human infrastructures and forest management plans need to be revisited.

Considering that demographic growth has increased over the last year in Canada (including in Québec), with the highest annual population growth rate (+2.7%) on record in 2022 since 1957 (population increase over 1 million people in 2023; Statistics Canada 2023) and will continue to increase in the future, more and more communities and infrastructures will be exposed to forest fires. In this respect, there is an urgent need for a risk zonation that considers multiple risks, including wildfires. Such a zonation will help municipalities and communities to better plan their future land use and development according to environmental hazards. Only taken into account the total costs of various insured properties from natural disasters in Québec (forest fires and floods being the most frequent and costly disasters), the first 9 months of 2023 were the most costly in the province, i.e., 612 M$ in 2023 compared to an average of around 97 M$ over the 5 years in 2011-2015, and around 222 M$ in 2016-2020 (CatIQ data, B. Marchand, personal communication). With the increase of economic costs from forest fires (Hope et al. 2016), this will induce either potential growth or potential closure in private insurance properties or industrial assets (ex., from forestry companies) in the future, making socioeconomic vulnerability an issue of concern for people and infrastructures located within the Intensive Protection Zone and abroad.

Other aspects that need to be considered or further studied are the cascade effects from extensive forest fires, notably on water cycle and on the occurrence of floods from runoff and sediment or material transport that will potentially increase from burned areas within watersheds (Robinne et al. 2018). Note that watershed morphology and forest stand density interact with large floods, through large wood and organic matter jams (Lininger et al. 2021). Forest fires will also affect the seasonality of floods in spring months, during rain on snow events, but also in summer and fall months during intense or extreme precipitation events, that are expected to increase over the following years (IPCC 2023). Not to mention the amplification of mercury concentrations in fish throughout the watershed (Garcia and Carignan 2005), which can have an impact on the health of indigenous communities living off the land. All these domino effects and potential damaging tipping points are important reasons for concern and need to be integrated in risk management approaches.

The increase in systemic risks, from various hydrometeorological hazards or extensive risks from regular and combined ones occurring at the regional or local scale, will set a clear limit to current adaptation options (O’Neill et al. 2017). Integrated risk management with a holistic overview of various climate risks could be an asset, with different strategies to propose from individual to community-centered approaches, in putting forward the advantages from specific interventions to the most vulnerable communities and forest ecosystems, and in considering the reduction of forest fires consequences.

## Conclusions

After the 2023 extreme fire season, Québec saw concerted efforts among various stakeholders and First Nations (e.g., Sommet sur les feux de forêt [ORRFB 2023]) to challenge the untenable *status quo* and the need for change considering an extreme fire season and ongoing climate changes. A consultation on forest management was initiated by the Québec Minister of Natural Resources and Forests on 15 November 2023, with a view to adapt to this new reality. Current forest management strategies may no longer be sufficient, requiring a revision of strategies related to the sustainable management of forests, including wood production, protected areas, and wildlife habitats. Such reflections offer an opportunity to update operational methodologies with sustainability principles, particularly for the forest sector, communities, First Nations, and ecosystems.

Our analysis emphasizes that it is crucial to put the impacts and consequences of the 2023 wildfires in the context of increasing fire activity due to climate change. A continued rise in wildfires under business-as-usual practices could lead to a decline in boreal forest health and its socio-ecological services, impacting the forest industry, carbon sequestration, wildlife and their habitat, and cultural values for indigenous and non-indigenous communities. Risk assessment and mitigation, along with adaptation and rapid actions, are key. This includes redefining forestry to make ecosystems and the forest sector more resistant and resilient, identifying vulnerabilities and co-benefits, and implementing regional adaptation measures integrating diverse expertise (Boulanger et al. 2023). Costs of adaptation strategies must be considered, prioritizing approaches with multiple mutual benefits. A precautionary approach is crucial in the face of uncertain warming (Millar et al. 2007). Systemic risks of forest fires and climate change impacts demand an integrated risk management approach, enhancing preventive tools and early-warning systems. Experiences like the one we faced during the 2023 fire season are opportunities to remind ourselves that we need to improve fire and forest management strategies and policies. This holistic approach would enhance our ability to predict, prevent, and respond to forest fires, reducing their impacts on economic sectors, ecosystems, and people.

## Appendices

### Supplementary Material S1 - Weather parameters

**Figure S1.1.**
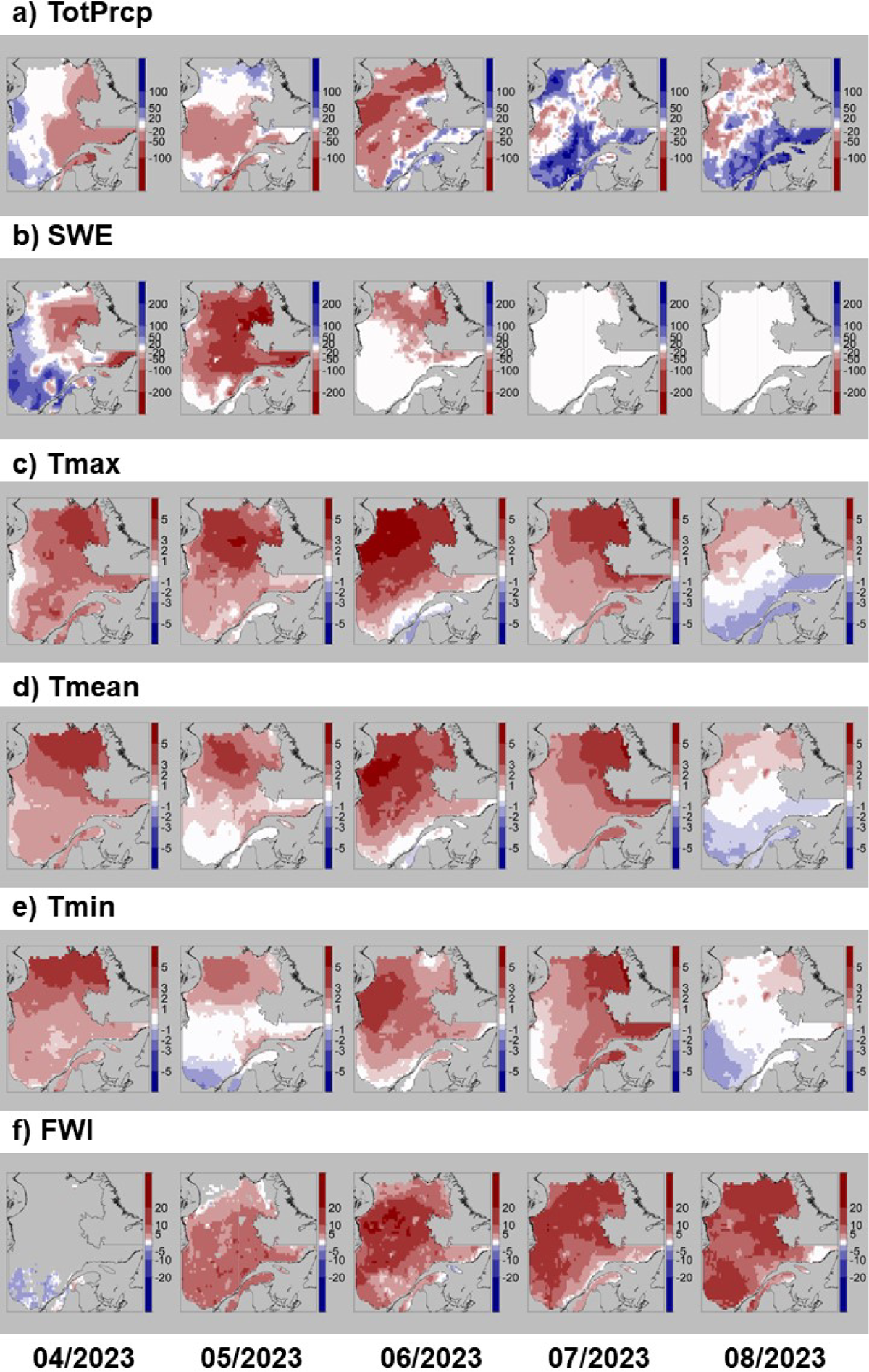
Mapped monthly anomalies from April to August 2023 relative to the 1991-2010 for a) total precipitation (TotPrcp, in mm), b) snow water equivalent (SWE, in mm), c) maximum daily temperature (Tmax, in celcius), d) mean daily temperature (Tmean, in celcius), e) minimum daily temperature (Tmin, in celcius) and f) fire-weather index (FWI, unitless). Values were assessed using ERA reanalyses (Hersbach et al. 2020)

**Figure S1.2.**
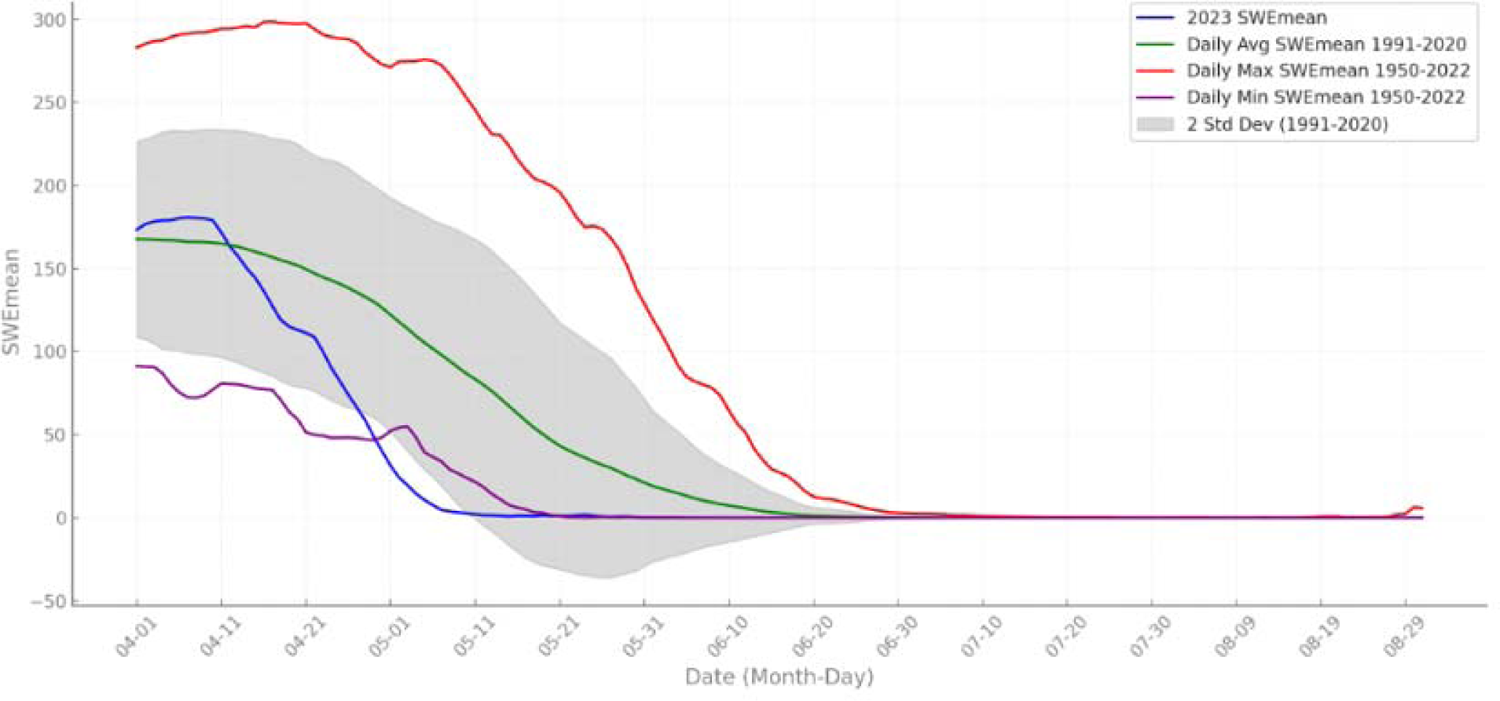
Daily values for snow weather equivalent (SWE) from April 1st to August 31st. Values for 2023 (blue line) against the historical average (1991-2020, green line) and the historical daily range (maximum in red, minimum in purple) for the period 1950-2022. A gray shading around the average indicates the range of two standard deviations using the 1991-2020 daily data. PrcpTot and DSR values are represented as cumulative sums starting from April 1st. values were computed for the area covered by homogeneous fire regime zones in Québec.

**Figure S1.3.**
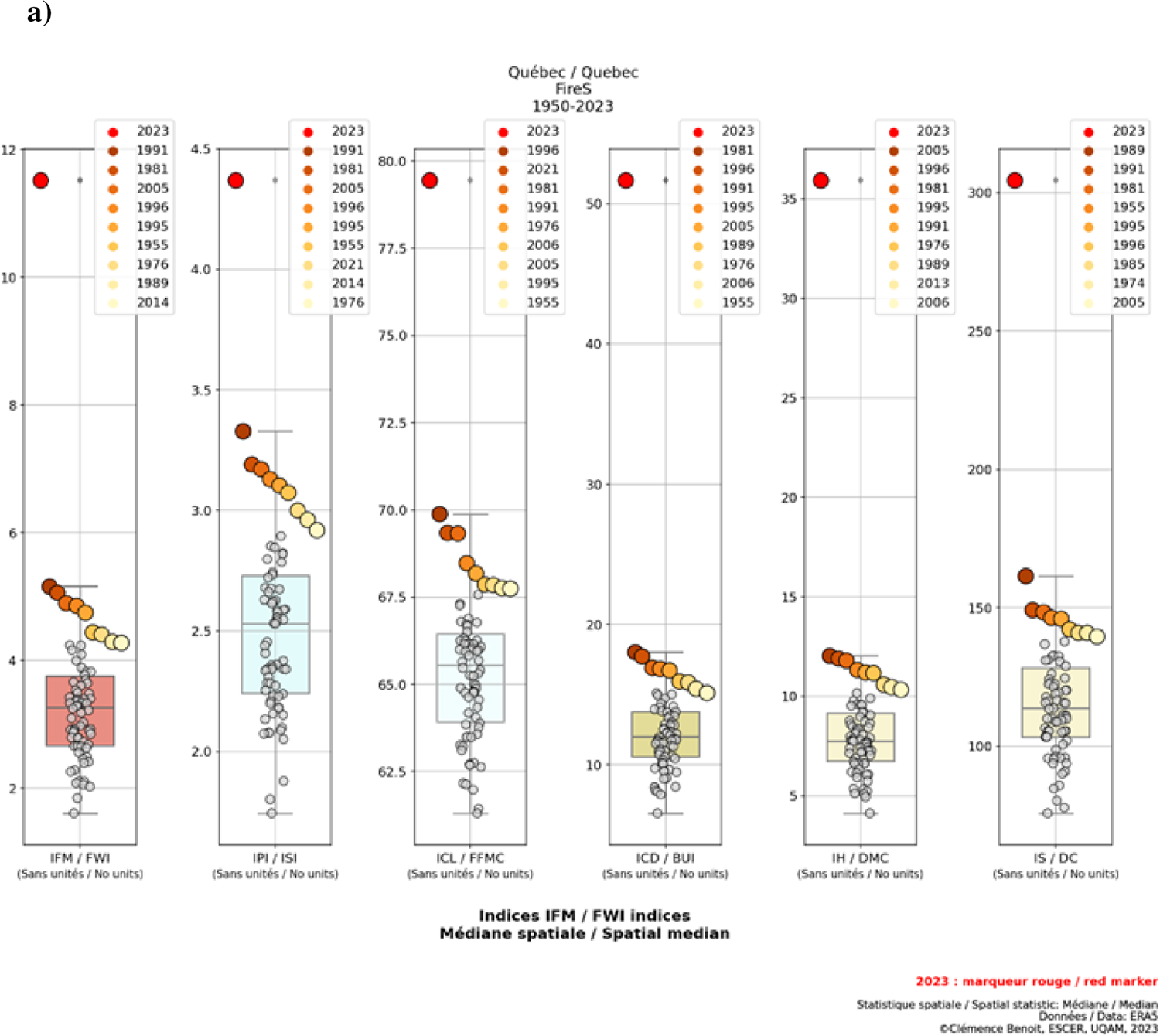
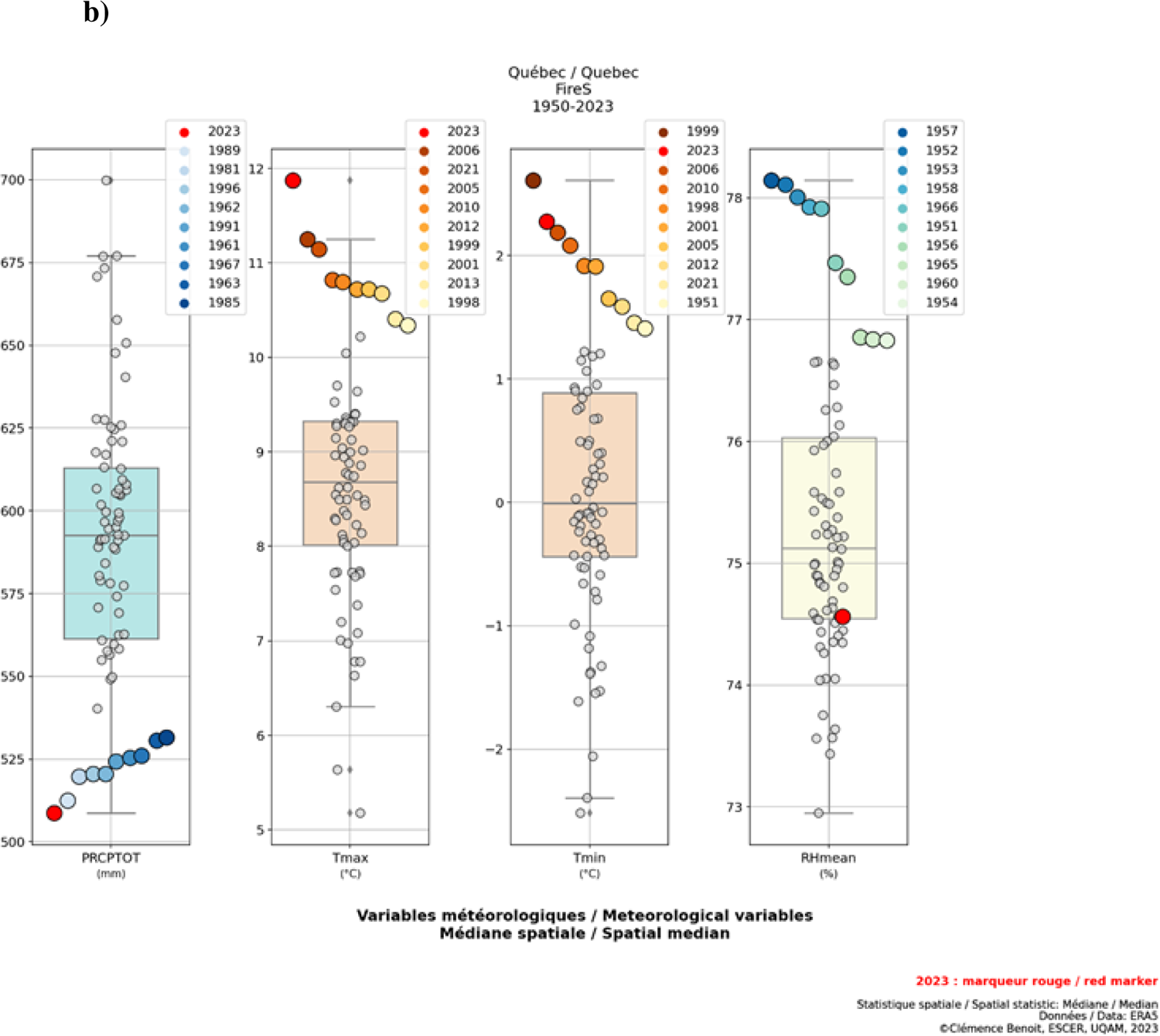
Boxplots of FWI indices (a) and meteorological variables (b) for all years from 1950 to 2023 during the fire season (May to October) for all of Québec. The year 2023 is represented by a red circle. Raw data to compute FWI indices and meteorological variables are from the ERA5 reanalysis (Hersbach et al. 2020).

**Figure S1.4.**
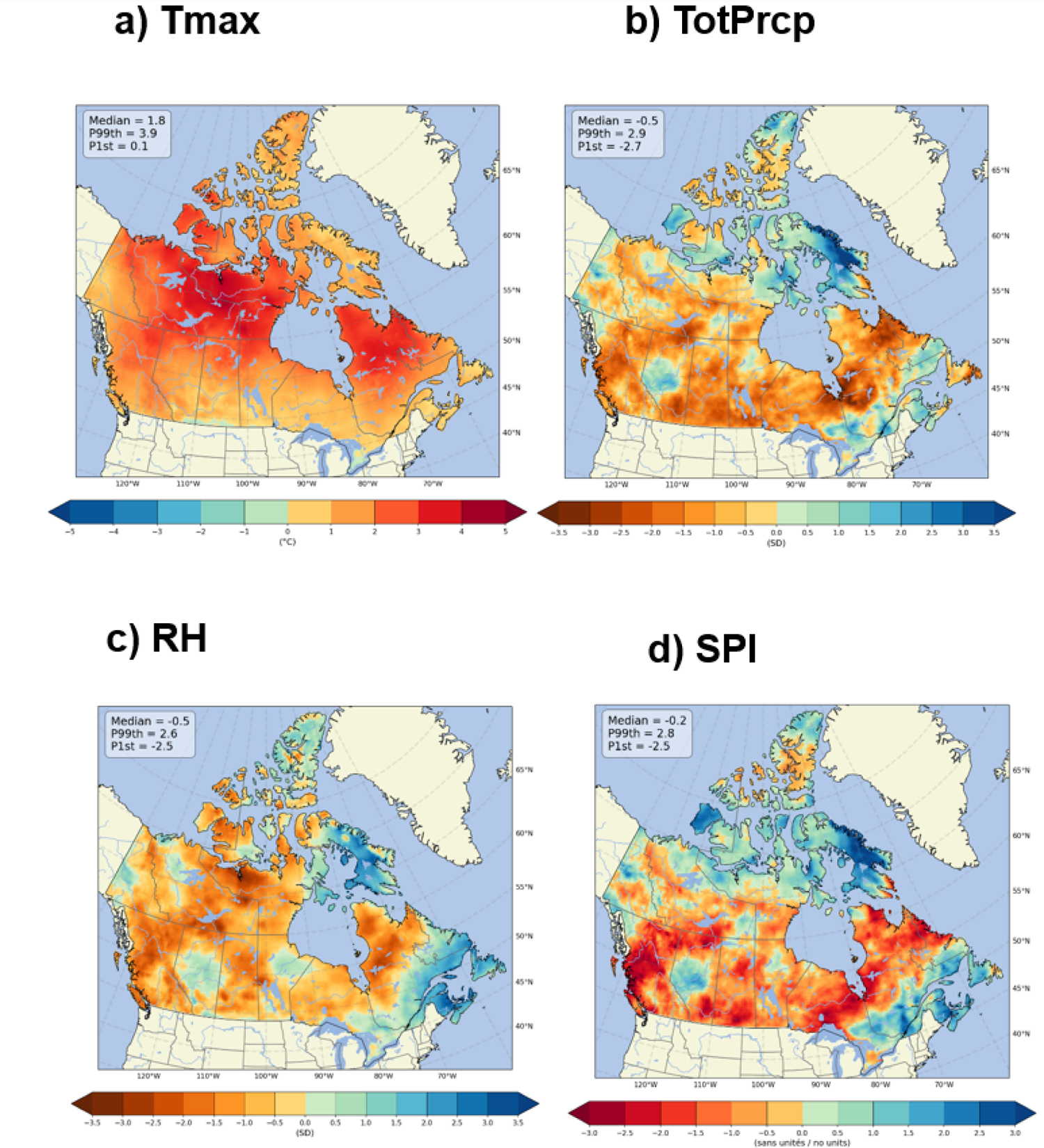
Anomalies over the entire 2023 fire season (May to October inclusive) of: a) daily maximum temperatures (Tmax, in °C), b) cumulative total precipitation (TotPrcp, unitless, standardized values), and c) relative humidity (RH, unitless, standardized values). d) Standardized Precipitation Index values calculated over 6 months from May to October in 2023 (SPI, −2< extremely dry and > 2 extremely wet). All anomalies are calculated relative to climatology from 1991 to 2020. Data are from the ERA5 reanalysis (Hersbach et al. 2020).

## Supplementary Material S2. - Climatic Data for Historical Drought Code Analysis

We collected daily maximum temperature (°C) and precipitation (mm) data from January 1, 1900, to October 23, 2023, using BioSIM v11 software. This software utilizes site-specific estimates derived from Environment and Climate Change Canada’s historical daily weather observations, which are available at ftp://ftp.cfl.forestry.ca/regniere/Data/Weather/Daily/ (last accessed on 2023-10-23). The daily weather data were interpolated onto a 0.5° x 0.5° grid, based on the four nearest weather stations, using inverse distance weighting. This interpolation accounted for variations in elevation and location, adjusting for regional gradients (Régnière et al., 2014).

Utilizing daily maximum temperature and precipitation data, we applied the Fire Weather Index System module within the BioSIM software to calculate the daily Drought Code component for each grid point. This approach follows the methodology outlined by Terrier et al. (2013) and Gaboriau et al. (2022). We assumed the start of the fire season to be on the third day after snowmelt or on the third consecutive day with a noon temperature above 12°C, whichever came first (Lawson and Armitage, 2008; coefficients *a* and *b* were set to 0.5). We also considered the overwintering effect, which represents the previous year’s legacy index values at the end of the fire season, rather than simply resetting the index computation. Similarly, we assumed the fire season ended with the accumulation of 2 cm of snow on the ground for seven consecutive days or after three consecutive days with a daily minimum temperature below 0°C, whichever occurred first (Lawson and Armitage, 2008). BioSIM generated snow depth information using a snowpack model driven by air temperature and precipitation values, as described by Brown et al. (2003).

For each grid point, we ranked the severity of the Drought Code (DC) for each day in 2023 against the historical DC severity for the same date over a 124-year period. Daily maps illustrating these rankings were then created. To generate these maps, we applied an inverse distance weighting algorithm to the gridded point locations, resulting in a continuous raster product covering eastern boreal forests.

Next, for each grid point, we calculated monthly DC time series by averaging the daily DC values over June. We further averaged these series within homogeneous fire regime zones and analyzed changes in the occurrence rates of years with extreme monthly June DC severities using the procedure outlined by Girardin et al. (2009). Extreme June DC severity was defined as years in which June DC exceeded a detection threshold computed from a running median smoothing (2k+1 points) and the median of absolute distances to the median (factor z) computed over the period 1900-2023. This approach is particularly valuable when both the background state and interannual variability change over time. Kernel estimation allows for a detailed examination of time-dependent event occurrence rates and an assessment of significant changes with the help of confidence bands. The running median smoothing parameter k was set to 40 years, and the number of extreme events under analysis was set to approximate the highest 15th percentile (a function of the distance to the median z, varying by increments of 0.5). The kernel bandwidth, h, was set to 15 years. Confidence bands (90%) around λ(t) were determined using a bootstrap technique. These confidence bands facilitated the assessment of whether peaks and troughs in the occurrence of extreme events were statistically significant.

## Supplementary material S3 - Potential impacts of the 2023 wildfires on timber supply in Québec: an example in UG 107

### Impact of wildfire on wood supply: simulation details

Wood supply is based on two main components: the sustained yield cutting rate, which depends largely on the average commercial maturity age, and the productive area included in the annual allowable cut (AAC) calculation. Forest fires influence both these components. In the short term, by consuming mature forests, they have a direct effect on the volumes available for harvesting. In the long term, by generating regeneration failures, they reduce productive areas.

This section analyzes these two major impacts on wood supply as calculated by the *Bureau du Forestier en Chef* (BFEC): the volume loss of commercially mature wood and the loss of productive area due to regeneration failures.

### Losses in mature timber volumes

The assessments carried out are based on the analysis of one management unit (UG) particularly affected by the fires of spring 2023, i.e., Unit 107 (see figure S4.1), which lies to the east of Lebel-sur-Quévillon. The forest in this unit covers almost 1.3 million ha and the fires spread over 281,818 ha, or 21.8% of the forested area in 2023. If we consider only the mature portion of the forest, nearly 26% of it burned. What are the impacts of these losses on the AAC?

In Québec, the AAC calculation i.e., the calculation of the maximum volume that can be allocated to harvesting, taking into account a set of constraints limiting the areas included in the calculation or forest yields, is revised every 5 years. It is during these revisions that losses caused by forest fires are considered. This *a posteriori* approach to fire risk management is appropriate for low-risk areas. If we exclude the year 2023, we can say that the Nord-du-Québec region, which includes UG 107, was at low risk, with an average yearly burn rate of 0.32% for the period 1972-2022.

However, this burn rate is well below historical rates, which range from 0.45% to 0.83% depending on the area (Couillard et al. 2022). During the period 1972-2021, a total of 15% of forest area in the Nord-du-Québec region was burned. In comparison, 12% of the region’s forest area was burned during spring 2023. The 2023 season thus reminds us that we are indeed in a high-risk area according to historical data covering the period 1890-2020 (Couillard et al. 2022), even though the last 50 years have been little affected by fires.

Forest fire impacts on UG 107’s AAC is likely to be major, as the allowable cut calculations were established in the absence of consideration for fires. Additionally, the volume loss available for harvesting as a result of fires will translate almost directly into a loss of AAC. It is possible to mitigate these losses by speeding up the salvage logging, but even so, the industry estimates that it could only recover 10 to 15% of burnt wood volumes (Le Quotidien 2023), as the recovery window is very short due to wood degradation caused by long horned beetles.

### How would a wood buffer have limited the impact of wildfire on AAC?

To answer this question, we simulated (using a model developed in Leduc *et al*. 2015) how a wood buffer established 20 years ago would have limited the current decline in AAC. To do this, we first calculated the rejuvenation rates of the mature forest (over 80 years old) between 2002 and 2022 (Figure S3.1). This rate essentially reflects the harvesting rate, but also includes the burning rate. This rejuvenation rate needs to be adjusted according to the productivity of the forests. For UG 107, the latest BFEC estimates put the time required to produce a mature stand of black spruce at 90 years, implying an overall rejuvenation rate of no more than 1.1%. In UG 107 for the period 2002-2022, by comparing mature areas in 2002 with those in 2022, this rejuvenation rate is estimated at 1.39% per year on average, equivalent to an average harvest rotation of 72 years. By way of comparison, the current harvest rate reviewed by the BFEC for the 2023-28 period is 1%.

In our simulation, we then reduced the average 2002-2022 rate to 1.11% and 1.25%, corresponding to a buffer of 20% and 10%. We took the 2002 age structure, applied the reduced harvesting rate over a 20-year period and then burned the forest with the same areas recorded in 2023. We then obtained the area after fire in 2023 for each age class, and thus the residual unburnt area in mature forest. We then applied the current harvesting rate to calculate the number of harvesting years available in each scenario. This number of years is compared with the time required for immature stands to reach commercial maturity. If the number of harvesting years is greater than the time required to refill mature stands, we conclude that the harvesting rate is sustainable. However, if the number of years of harvesting appears to be less than the maturation period, the harvesting rate must be reduced in order to increase the number of years of harvesting. For the scenarios under study, without precautionary reserves but including salvage logging, we estimate that the reduction will be 20-22% for UG 107. Thus, a 20% buffer established in 2002, would have ensured that there would be no reduction in AAC following the fires of 2023 (Figure S3.2). However, the 10% reserve generates a slight shortfall, since the maturation period is 29 years, whereas the number of years of operation is 27. This difference should result in a slight drop in AAC of between 5% and 8%. Note that this reduction will be felt over several decades of exploitation, as the forest is very slow to regrow. This is important when estimating the cost of setting up a precautionary buffer approach. Moreover, we note that having set a precautionary reserve in 2002 would result in more wood harvested, cumulatively, after 2040 in this area, when compared with a scenario where no precautionary reserve is applied (Figure S3.3).

### Loss of productive area due to regeneration failure

In natural conditions boreal forest regenerates after fires thanks to the aerial seed bank, which is released by the action of heat on the serotinous cones. In some cases, however, fire can lead to forest degradation if the stand is immature and has not had time to build up an aerial seed bank. In UG 107, almost 50% of the areas burnt during the 2023 fires were less than 50 years old and may need to be reforested. Of these areas of young forest burnt, 66% were cutover and 23% reforested. In fact, although fire is the proximal factor causing regeneration failure, the clearcutting that has significantly rejuvenated the boreal forest in recent decades is a predisposing factor. In fact, before the fires of 2023, the age structure of UG 107 was estimated to contain twice as many young stands as the historical proportion for the same area. Today, 67% of these stands are at risk of regeneration failure because they are less than 50 years old (figure S3.1).

We estimate that the overall effect of fire is the result of short-term effect on the volumes available for harvesting, plus long-term effect on the productive areas included in the calculation. For UG 107, our estimates indicate that the decline resulting from the loss of available harvest volume will be 20-22%. To this must be added a 10% quasi-permanent loss of productive area, for a total of 32% reduction in allowable cut.

**Figure S3.1.**
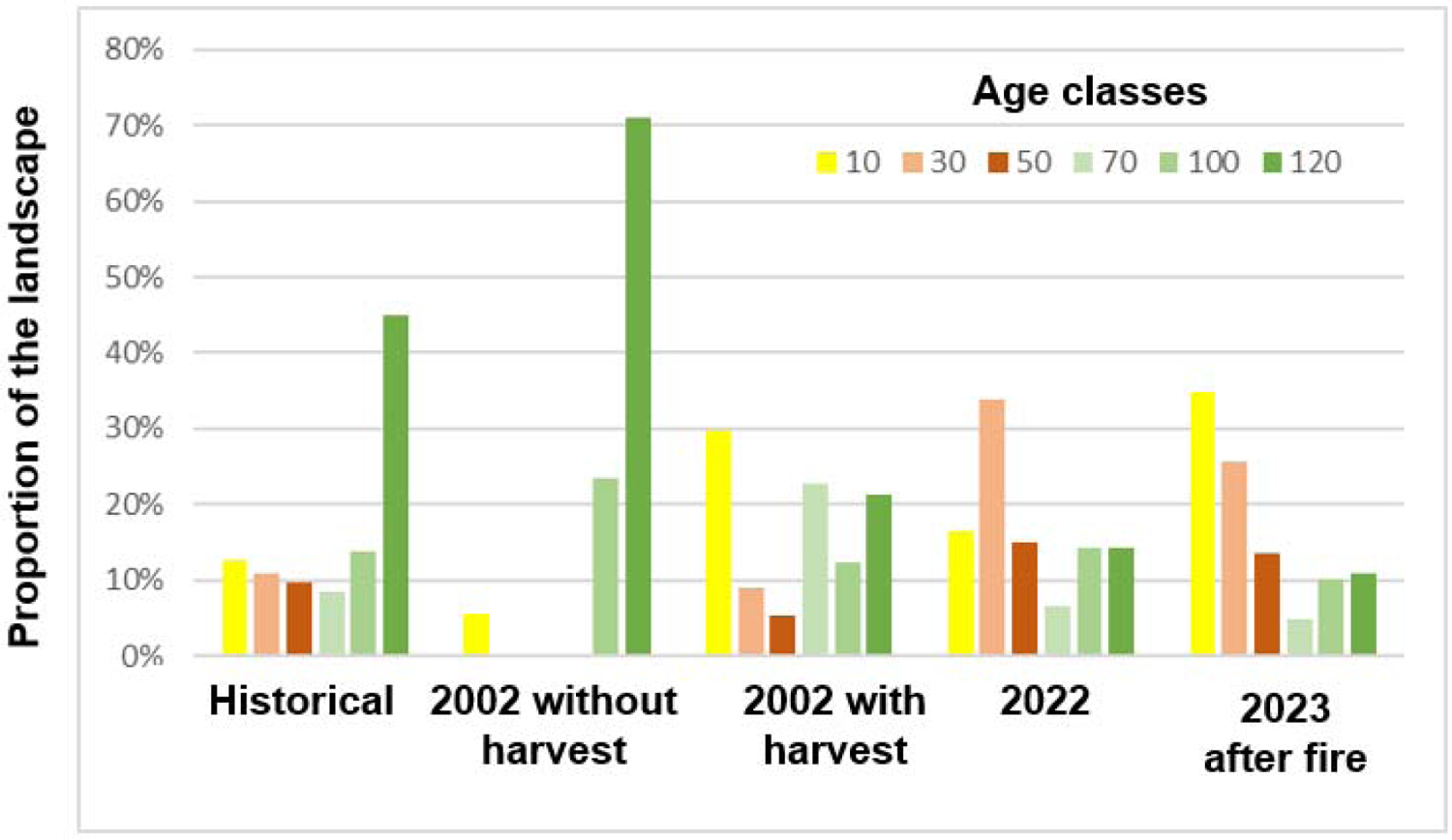
Age structure of UG 107 at different times. The historical age structure is deduced from an average fire cycle of 150 years. The age structure for 2002 without logging reflects the virtual absence of burnt areas during the period 1940-2002. The 2002 age structure with harvesting was produced using the 3rd inventory ecoforestry map. The 2022 age structure was calculated using the 4th inventory forest map. The 2023 age structure was produced using Sentinel post-fire satellite images, which precisely delineate the burnt areas.

**Figure S3.2.**
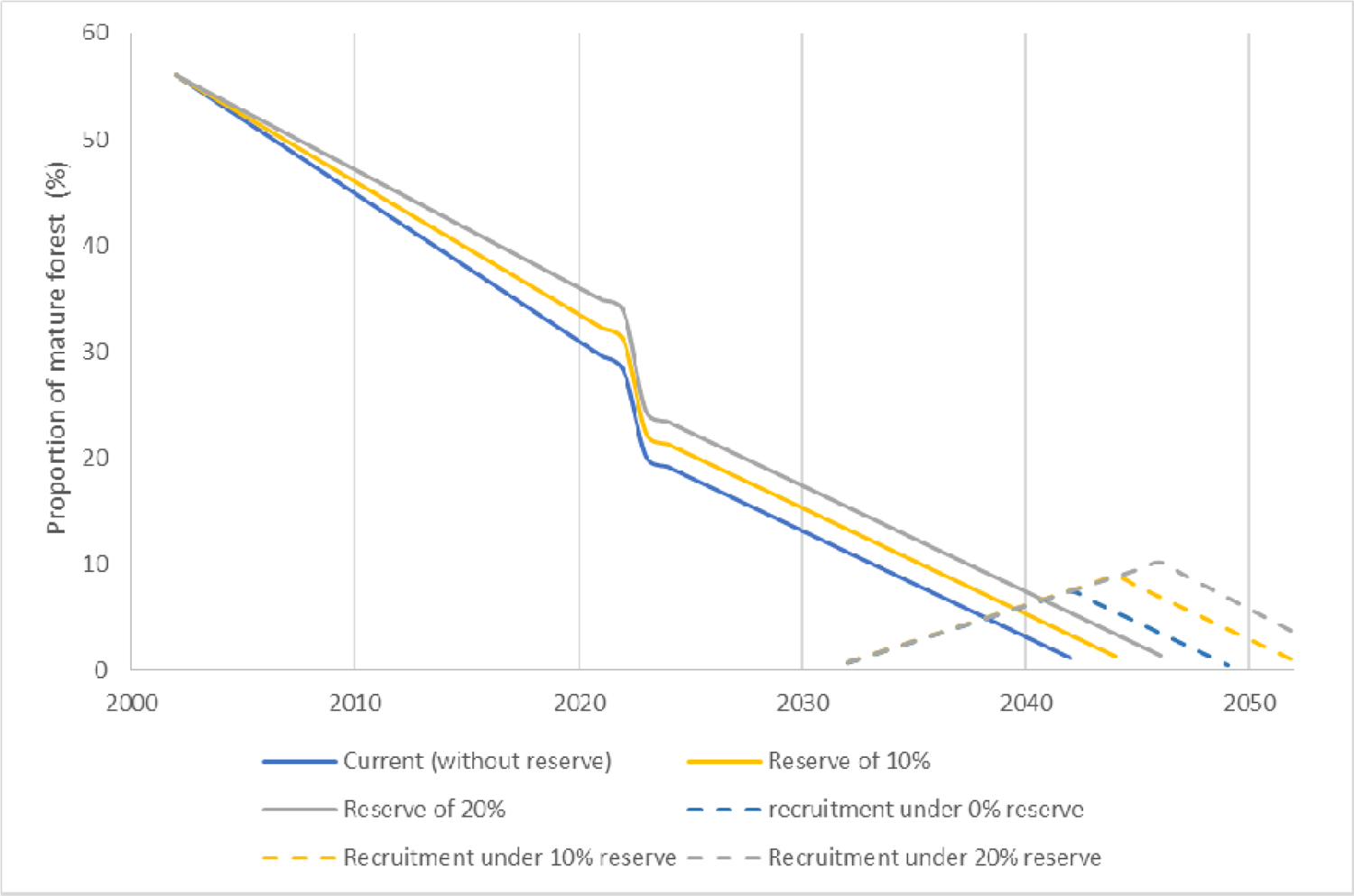
Evolution of forest area available for harvesting in UG 107 according to 3 scenarios: a) current scenario with no wood reserve (blue line); with a 10% reserve implemented in 2002 (yellow line); with a 20% reserve (gray line). Harvesting rates before 2023 are in line with those observed during the 2002-2022 period. From 2023 onwards, those rates are reduced to 1%, i.e. to levels updated for the period 2023-2028. The sudden drop in 2023 reflects losses due to fires. The dashed lines indicate recruitment due to the maturation of 40-60 year-old stands after the 2023 fire. With no post-fire decline in AAC, the scenario with no wood reserve depletes all mature areas in 2049, i.e. 4 years before 2052, when recruitment would be ensured by the stands that were 20-40 year-old stands in 2002, at the time when reserves could have been established.

**Figure S3.3.**
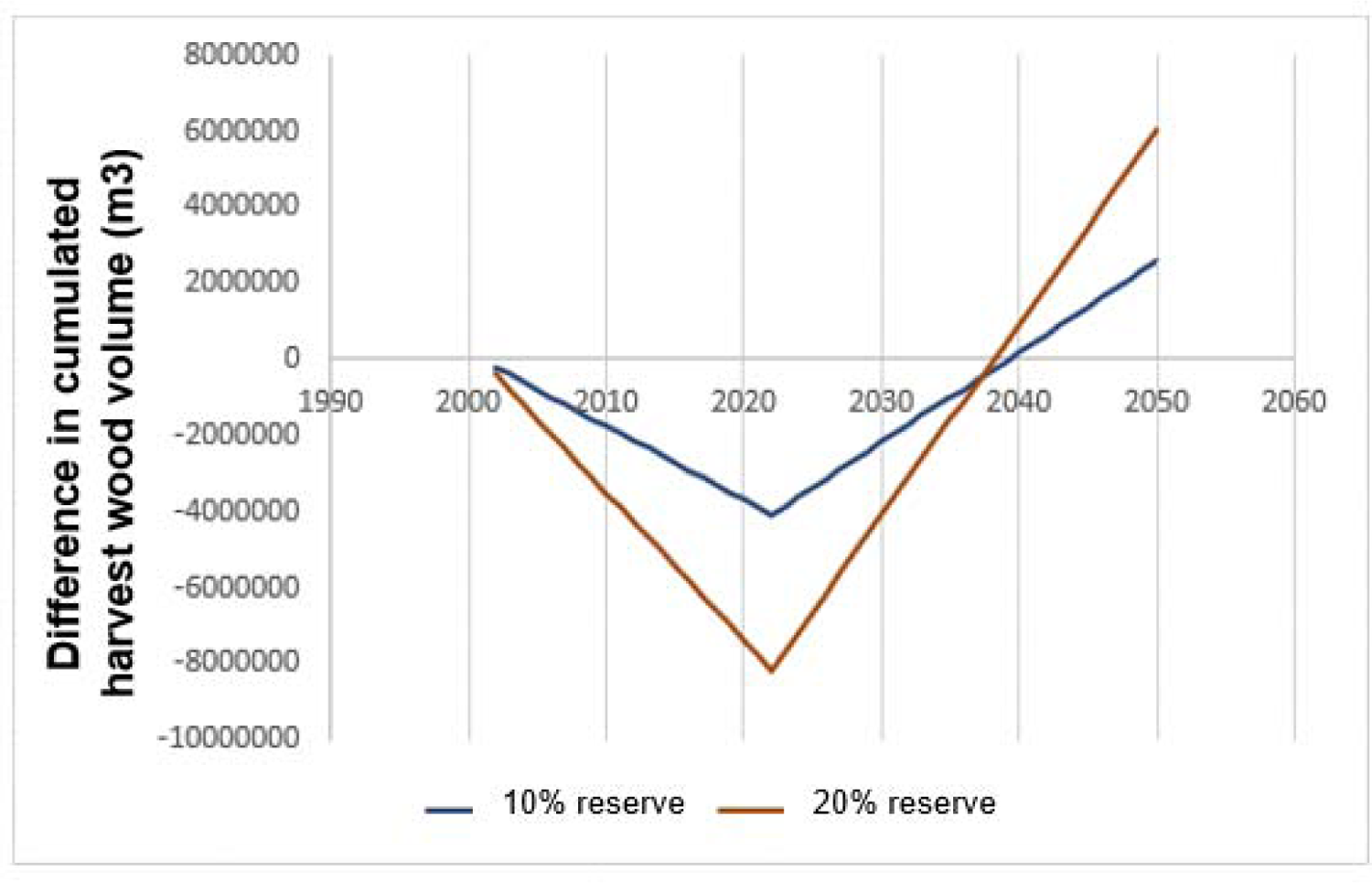
Differences in cumulated harvested volumes between scenarios where a precautionary reserve of 10% (blue) or 20% (red) is adopted in 2002 compared with a scenario where no precautionary reserve is adopted. The trough corresponds to the 2023 fires. Even though the volume harvested in the first years is less when a precautionary reserve is used, this approach leads to greater overall volume harvested after around 2040, compared to a method that does not include a precautionary reserve.

## Supplementary material S4 - Detailed assessment of what has burned in 2023 within Québec’s managed forests

This supplementary material presents a detailed assessment of the characteristics of Québec’s managed forests that have burned in 2023. We conducted such an assessment by combining different spatial data sources. We also present a preliminary analysis of the selectivity of fire towards certain forest stand types.

### Mapping burned area at medium resolution

Using the coarse polygons produced by Québec’s *Société de Protection des Forêts Contre le Feu* (SOPFEU), we first identified sectors affected by fire. These polygons are derived from coarse-resolution remote sensed data (https://firms.modaps.eosdis.nasa.gov) and thus cannot be used for our assessment objectives because they integrate many unburned forests. We thus used these polygons as a basis to build a high-resolution map of areas really affected by fire with a well-documented and widely used method: the Normalized Burn Ratio (NBR; see for ex.: Fernández-Manso et al. 2016; Boucher et al. 2017; Quintano et al. 2018 for more details). We extracted Sentinel-2 NBR pre- and post-fire 20m resolution rasters for each sector affected by fires. In the managed forests, fire activity stopped around June 27, 2023. Thus, all cloud-free Sentinel-2 images that were available between this date and August 15, 2023, were used for computing post-fire NBR rasters. Pre-fire NBR rasters were computed with all cloud-free Sentinel-2 available in the preceding year’s same period (July 27 to August 15, 2022). The differenced normalized burn ratio (dNBR) was then computed by subtracting 2022 to 2023 NBR values. Finally, areas affected by fire were defined by applying a threshold on dNBR values, which was validated with 600 burned and unburned points (visually defined on post-fire Short Wave Infrared RGB composite [SWIR] Sentinel-2 images).

**Figure S4.1.**
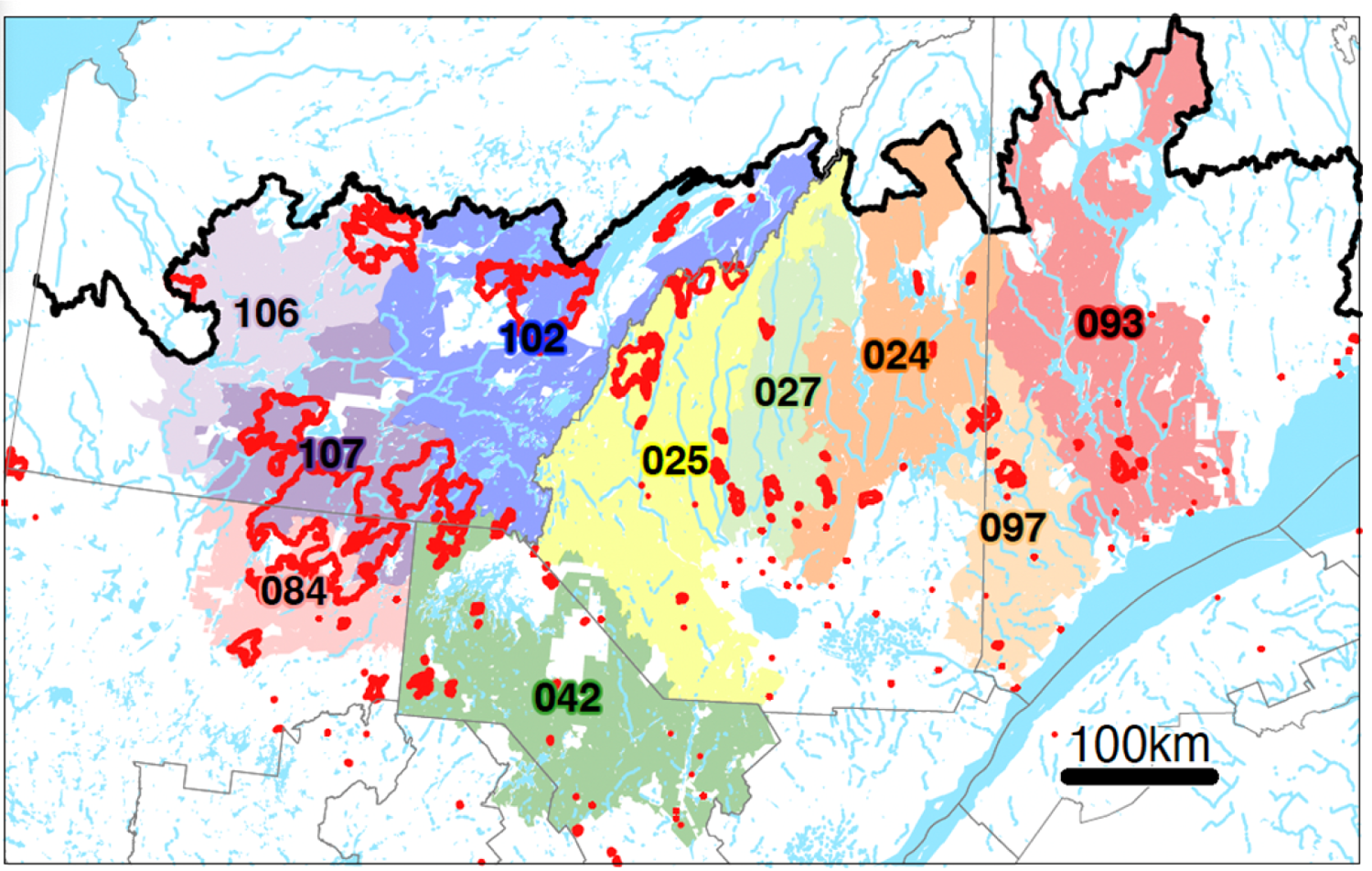
Index map of Québec’s forest Gestion Units (UG) that have been affected by fire in 2023 and for which assessment has been done for this study. A specific color represents each UG; their reference numbers are shown at their polygon centroids. Coarse SOPFEU perimeter of the 2023 fires are delimited in red. The thick black line in the upper part of the map shows the northern limit of managed forests, and fine grey lines show the boundaries of Québec’ administrative region.

### Assessing characteristics of forests that have burned in 2023

We then combined our medium-resolution fire maps with up-to-date Québec’s government forest maps (MFFP 2018; https://www.donneesQuebec.ca/recherche/dataset/carte-ecoforestiere-avec-perturbations). These maps contain 1:20,000 photo-interpreted polygons that estimate forest stand age and composition across the landscape. These maps also integrate information about non-forest site characteristics (e.g., peat bogs or bare rocky environments) and about the type of the last known disturbance (e.g., fire, logging). We first selected all burned areas classified as “productive forests” (i.e., managed forest sites), totalling 818 900 ha. We regrouped sites among those burned productive forests with three simplified classifications: age, composition, and pre-2023 disturbance type. Six simplified twenty-year age classes (0:20, 21:40, 41:60, 61:80, 81:100, >101 years old) were retained. The composition was grouped depending on tree species’ relative abundance among five large types: black spruce (>75% of black spruce), black spruce and jack pine (jack pine >25% and black spruce >25%, and totalizing > 75%), other or undifferenced conifers (conifers other than black spruce and jack pine [or undifferenced] >75%), mixed or hardwood (hardwood species >25%) and unknown (stands for which no information about composition is available in the forest maps). We also simplified information on pre-2023 disturbance types into four groups: logging, plantation, fire, and other/unknown. Analysis of the total area burned in each age, composition, and last disturbance type was first made for all of Québec’s managed forests affected by fires in 2023. Analyses were then subsequently made for each forest management Unit (UG; Figure S4.1) that has been significantly affected by fire in 2023. All results are presented as a summary table (Tables S4.1 to S4.22).

### Analyzing the selectivity of fire upon different forest characteristics

We also took advantage of these data to conduct a preliminary analysis of fire selectivity across different forest types. We compared the area of forest stand types that have been affected by fire (i.e., according to our medium-resolution burned maps) with the forest stands that were “available” within the landscape. Available forest stands within the landscape were defined as all forest stands within a 500-meter buffer around coarse-resolution SOPFEU polygons. We analyzed fire selectivity for forest stand types defined above (i.e., age, composition and pre-2023 disturbance) simply by comparing the proportion of a stand type’s total burn area (i.e., % affected by fire) to its proportion of total abundance within the landscape (i.e., % of availability). A forest stand type whose proportion of total burned area is less than its proportion of availability would thus be suspected of fire-resistant. In the opposite situation, the forest stand types would be considered fire-prone. The northernmost landscapes affected by the 2023 fires showed a very low heterogeneity in forest stand types (age, composition, and pre-2023 disturbance). We thus only retained the large fire complex of southwestern Québec (Figure S4.2) for those analyses because their heterogeneity in stand types is much more relevant to conducting such analyses.

**Table S4.1.** Burned areas by age class (columns) and composition groups (rows) for all the managed forests that burned in 2023 (i.e., productive forests south of the northern limit of managed forests). Results are expressed in 1000 ha (Kha) and % of the total.

|  | [0:20] |  | [21:40] |  | [41:60] |  | [61:80] |  | [81:100] |  | [101+] |  | Total |  |
| --- | --- | --- | --- | --- | --- | --- | --- | --- | --- | --- | --- | --- | --- | --- |
|  | Kha | % | Kha | % | Kha | % | Kha | % | Kha | % | Kha | % | Kha | % |
| Black spruce | 24.3 | 3 | 22.8 | 2.8 | 22.2 | 2.7 | 41.3 | 5 | 73.8 | 9 | 171.7 | 21 | 356.2 | 43.5 |
| Jack pine-Black sp. | 3.6 | 0.4 | 12.3 | 1.5 | 15.6 | 1.9 | 24.6 | 3 | 45.2 | 5.5 | 21.3 | 2.6 | 122.6 | 15 |
| Conifers (oth/undif) | 15 | 1.8 | 101.7 | 12.4 | 9.3 | 1.1 | 1.8 | 0.2 | 1.2 | 0.1 | 8.4 | 1 | 137.4 | 16.8 |
| Mixed-hardwood | 12.5 | 1.5 | 49.9 | 6.1 | 8.7 | 1.1 | 4.1 | 0.5 | 6.6 | 0.8 | 8.1 | 1 | 90 | 11 |
| Unknown | 71.9 | 8.8 | 36.8 | 4.5 | 3.9 | 0.5 | 0 | 0 | 0.1 | 0 | 0 | 0 | 112.7 | 13.8 |
| Total | 127.3 | 15.5 | 223.6 | 27.3 | 59.7 | 7.3 | 71.8 | 8.8 | 126.9 | 15.5 | 209.6 | 25.6 | 818.9 | 100 |

**Table S4.2.** Burned areas by age class (columns) and pre-2023 disturbance types (rows) for the managed forests that burned in 2023 (i.e., productive forests south of the northern limit of managed forests). Results are expressed in 1000 ha (Kha) and % of the total.

|  | [0:20] |  | [21:40] |  | [41:60] |  | [61:80] |  | [81:100] |  | [101+] |  | Total |  |
| --- | --- | --- | --- | --- | --- | --- | --- | --- | --- | --- | --- | --- | --- | --- |
|  | Kha | % | Kha | % | Kha | % | Kha | % | Kha | % | Kha | % | Kha | % |
| Logging | 60.9 | 7.4 | 125.8 | 15.4 | 15.5 | 1.9 | 3.8 | 0.5 | 0 | 0 | 0 | 0 | 206.1 | 25.2 |
| Plantation | 34.3 | 4.2 | 46 | 5.6 | 0.2 | 0 | 0 | 0 | 0 | 0 | 0 | 0 | 80.5 | 9.8 |
| Fire | 19.2 | 2.3 | 34.2 | 4.2 | 17.1 | 2.1 | 9.8 | 1.2 | 27 | 3.3 | 13.2 | 1.6 | 120.4 | 14.7 |
| Other/unknown | 12.8 | 1.6 | 17.6 | 2.1 | 26.9 | 3.3 | 58.2 | 7.1 | 100 | 12.2 | 196.3 | 24 | 411.8 | 50.3 |
| Total | 127.3 | 15.5 | 223.6 | 27.3 | 59.7 | 7.3 | 71.8 | 8.8 | 126.9 | 15.5 | 209.6 | 25.6 | 818.9 | 100 |

**Table S4.3.** Burned areas by age class (columns) and composition groups (rows) for productive forests in the **UG 106**. Results are expressed in 1000 ha (Kha) and % of the total.

|  | [0:20] |  | [21:40] |  | [41:60] |  | [61:80] |  | [81:100] |  | [101+] |  | Total |  |
| --- | --- | --- | --- | --- | --- | --- | --- | --- | --- | --- | --- | --- | --- | --- |
|  | Kha | % | Kha | % | Kha | % | Kha | % | Kha | % | Kha | % | Kha | % |
| Black spruce | 0.6 | 0.9 | 0.7 | 1.1 | 0.8 | 1.3 | 1.3 | 1.9 | 3.7 | 5.5 | 27.7 | 41.4 | 34.9 | 52.1 |
| Jack pine-Black sp. | 0.1 | 0.1 | 0.8 | 1.2 | 0.8 | 1.2 | 2.5 | 3.8 | 3.5 | 5.2 | 4.2 | 6.2 | 11.8 | 17.6 |
| Conifers (oth/undif) | 0.4 | 0.5 | 0.7 | 1.1 | 0.2 | 0.2 | 0.1 | 0.1 | 0.1 | 0.2 | 0.2 | 0.3 | 1.6 | 2.4 |
| Mixed-hardwood | 0.3 | 0.4 | 3 | 4.5 | 0.3 | 0.5 | 0.1 | 0.1 | 0.2 | 0.4 | 0.4 | 0.6 | 4.4 | 6.5 |
| Unknown | 9.3 | 13.8 | 4.9 | 7.3 | 0.2 | 0.3 | 0 | 0 | 0 | 0 | 0 | 0 | 14.3 | 21.4 |
| Total | 10.6 | 15.8 | 10.2 | 15.2 | 2.3 | 3.4 | 4 | 5.9 | 7.5 | 11.1 | 32.5 | 48.5 | 67 | 100 |

**Table S4.4.** Burned areas by age class (columns) and pre-2023 disturbance types (rows) for productive forests in the **UG 106**. Results are expressed in 1000 ha (Kha) and % of the total.

|  | [0:20] |  | [21:40] |  | [41:60] |  | [61:80] |  | [81:100] |  | [101+] |  | Total |  |
| --- | --- | --- | --- | --- | --- | --- | --- | --- | --- | --- | --- | --- | --- | --- |
|  | Kha | % | Kha | % | Kha | % | Kha | % | Kha | % | Kha | % | Kha | % |
| Logging | 5.3 | 7.9 | 0.4 | 0.7 | 0 | 0 | 0 | 0 | 0 | 0 | 0 | 0 | 5.7 | 8.6 |
| Plantation | 0.1 | 0.1 | 0 | 0 | 0 | 0 | 0 | 0 | 0 | 0 | 0 | 0 | 0.1 | 0.1 |
| Fire | 4.6 | 6.8 | 8.6 | 12.9 | 1.1 | 1.6 | 0.6 | 0.9 | 2.2 | 3.3 | 2.7 | 4 | 19.8 | 29.5 |
| Other/unknown | 0.7 | 1 | 1.1 | 1.6 | 1.2 | 1.8 | 3.4 | 5 | 5.2 | 7.8 | 29.8 | 44.5 | 41.4 | 61.8 |
| Total | 10.6 | 15.8 | 10.2 | 15.2 | 2.3 | 3.4 | 4 | 5.9 | 7.5 | 11.1 | 32.5 | 48.5 | 67 | 100 |

**Table S4.5.** Burned areas by age class (columns) and composition groups (rows) for productive forests in the **UG 107**. Results are expressed in 1000 ha (Kha) and % of the total.

|  | [0:20] |  | [21:40] |  | [41:60] |  | [61:80] |  | [81:100] |  | [101+] |  | Total |  |
| --- | --- | --- | --- | --- | --- | --- | --- | --- | --- | --- | --- | --- | --- | --- |
|  | Kha | % | Kha | % | Kha | % | Kha | % | Kha | % | Kha | % | Kha | % |
| Black spruce | 9.6 | 3.5 | 9.2 | 3.4 | 8.8 | 3.3 | 13.9 | 5.2 | 31.6 | 11.7 | 33.5 | 12.4 | 106.7 | 39.5 |
| Jack pine-Black sp. | 1.4 | 0.5 | 6 | 2.2 | 1.1 | 0.4 | 5.3 | 2 | 12.4 | 4.6 | 2.6 | 0.9 | 28.8 | 10.7 |
| Conifers (oth/undif) | 4.7 | 1.8 | 48.5 | 18 | 3.2 | 1.2 | 0.1 | 0 | 0.3 | 0.1 | 0.7 | 0.3 | 57.5 | 21.3 |
| Mixed-hardwood | 5.5 | 2 | 23.5 | 8.7 | 3.7 | 1.4 | 0.7 | 0.3 | 2.9 | 1.1 | 2 | 0.7 | 38.3 | 14.2 |
| Unknown | 29.2 | 10.8 | 9.3 | 3.4 | 0.1 | 0 | 0 | 0 | 0 | 0 | 0 | 0 | 38.6 | 14.3 |
| Total | 50.5 | 18.7 | 96.5 | 35.7 | 16.9 | 6.3 | 20.1 | 7.5 | 47.2 | 17.5 | 38.7 | 14.4 | 270 | 100 |

**Table S4.6.** Burned areas by age class (columns) and pre-2023 disturbance types (rows) for productive forests in the **UG 107**. Results are expressed in 1000 ha (Kha) and % of the total.

|  | [0:20] |  | [21:40] |  | [41:60] |  | [61:80] |  | [81:100] |  | [101+] |  | Total |  |
| --- | --- | --- | --- | --- | --- | --- | --- | --- | --- | --- | --- | --- | --- | --- |
|  | Kha | % | Kha | % | Kha | % | Kha | % | Kha | % | Kha | % | Kha | % |
| Logging | 23.7 | 8.8 | 63.1 | 23.4 | 4.5 | 1.7 | 0 | 0 | 0 | 0 | 0 | 0 | 91.2 | 33.8 |
| Plantation | 16.2 | 6 | 15.7 | 5.8 | 0 | 0 | 0 | 0 | 0 | 0 | 0 | 0 | 31.9 | 11.8 |
| Fire | 6.1 | 2.3 | 8.1 | 3 | 0.8 | 0.3 | 1.2 | 0.4 | 0 | 0 | 0.3 | 0.1 | 16.4 | 6.1 |
| Other/unknown | 4.5 | 1.7 | 9.6 | 3.6 | 11.7 | 4.3 | 18.9 | 7 | 47.2 | 17.5 | 38.4 | 14.2 | 130.4 | 48.3 |
| Total | 50.5 | 18.7 | 96.5 | 35.7 | 16.9 | 6.3 | 20.1 | 7.5 | 47.2 | 17.5 | 38.7 | 14.4 | 270 | 100 |

**Table S4.7.** Burned areas by age class (columns) and composition groups (rows) for productive forests in the **UG 084**. Results are expressed in 1000 ha (Kha) and % of the total.

|  | [0:20] |  | [21:40] |  | [41:60] |  | [61:80] |  | [81:100] |  | [101+] |  | Total |  |
| --- | --- | --- | --- | --- | --- | --- | --- | --- | --- | --- | --- | --- | --- | --- |
|  | Kha | % | Kha | % | Kha | % | Kha | % | Kha | % | Kha | % | Kha | % |
| Black spruce | 2.1 | 2.4 | 1.2 | 1.4 | 3.4 | 3.9 | 4.4 | 5 | 1.3 | 1.5 | 19.9 | 23 | 32.3 | 37.3 |
| Jack pine-Black sp. | 0.5 | 0.5 | 4.2 | 4.9 | 1.4 | 1.6 | 1.9 | 2.2 | 0.6 | 0.6 | 7.2 | 8.3 | 15.7 | 18.1 |
| Conifers (oth/undif) | 3.2 | 3.7 | 13.9 | 16 | 1.1 | 1.2 | 0.4 | 0.4 | 0 | 0 | 0.9 | 1 | 19.4 | 22.4 |
| Mixed-hardwood | 2.2 | 2.6 | 6.3 | 7.3 | 0.9 | 1.1 | 0.5 | 0.6 | 0.1 | 0.1 | 2.9 | 3.4 | 13 | 15 |
| Unknown | 5.2 | 6 | 1 | 1.1 | 0 | 0 | 0 | 0 | 0 | 0 | 0 | 0 | 6.2 | 7.1 |
| Total | 13.2 | 15.2 | 26.6 | 30.7 | 6.7 | 7.8 | 7.1 | 8.2 | 2 | 2.3 | 31 | 35.8 | 86.6 | 100 |

**Table S4.8.** Burned areas by age class (columns) and pre-2023 disturbance types (rows) for productive forests in the **UG 084**. Results are expressed in 1000 ha (Kha) and % of the total.

|  | [0:20] |  | [21:40] |  | [41:60] |  | [61:80] |  | [81:100] |  | [101+] |  | Total |  |
| --- | --- | --- | --- | --- | --- | --- | --- | --- | --- | --- | --- | --- | --- | --- |
|  | Kha | % | Kha | % | Kha | % | Kha | % | Kha | % | Kha | % | Kha | % |
| Logging | 6.8 | 7.9 | 14.1 | 16.3 | 4.6 | 5.3 | 0 | 0 | 0 | 0 | 0 | 0 | 25.5 | 29.5 |
| Plantation | 4.5 | 5.2 | 9.1 | 10.5 | 0.2 | 0.2 | 0 | 0 | 0 | 0 | 0 | 0 | 13.8 | 15.9 |
| Fire | 0.1 | 0.1 | 2.7 | 3.1 | 0.1 | 0.2 | 3.7 | 4.3 | 0.4 | 0.5 | 6.7 | 7.8 | 13.8 | 16 |
| Other/unknown | 1.8 | 2 | 0.6 | 0.7 | 1.9 | 2.1 | 3.4 | 4 | 1.6 | 1.8 | 24.2 | 28 | 33.5 | 38.7 |
| Total | 13.2 | 15.2 | 26.6 | 30.7 | 6.7 | 7.8 | 7.1 | 8.2 | 2 | 2.3 | 31 | 35.8 | 86.6 | 100 |

**Table S4.9.** Burned areas by age class (columns) and composition groups (rows) for productive forests in the **UG 102**. Results are expressed in 1000 ha (Kha) and % of the total.

|  | [0:20] |  | [21:40] |  | [41:60] |  | [61:80] |  | [81:100] |  | [101+] |  | Total |  |
| --- | --- | --- | --- | --- | --- | --- | --- | --- | --- | --- | --- | --- | --- | --- |
|  | Kha | % | Kha | % | Kha | % | Kha | % | Kha | % | Kha | % | Kha | % |
| Black spruce | 5.5 | 3.4 | 4.4 | 2.7 | 1.9 | 1.2 | 4.9 | 3 | 20.3 | 12.5 | 28.8 | 17.8 | 65.9 | 40.7 |
| Jack pine-Black sp. | 0.5 | 0.3 | 0.1 | 0.1 | 6.6 | 4.1 | 5.1 | 3.2 | 18 | 11.2 | 1.5 | 0.9 | 31.9 | 19.7 |
| Conifers (oth/undif) | 1.5 | 0.9 | 24.2 | 15 | 1.8 | 1.1 | 0.1 | 0.1 | 0.2 | 0.1 | 0.5 | 0.3 | 28.4 | 17.6 |
| Mixed-hardwood | 0.9 | 0.5 | 9.2 | 5.7 | 0.8 | 0.5 | 0.5 | 0.3 | 1.8 | 1.1 | 0.5 | 0.3 | 13.7 | 8.5 |
| Unknown | 11.9 | 7.4 | 7.5 | 4.6 | 2.4 | 1.5 | 0 | 0 | 0 | 0 | 0 | 0 | 21.9 | 13.5 |
| Total | 20.4 | 12.6 | 45.5 | 28.1 | 13.5 | 8.4 | 10.6 | 6.5 | 40.4 | 25 | 31.3 | 19.4 | 161.8 | 100 |

**Table S4.10.**
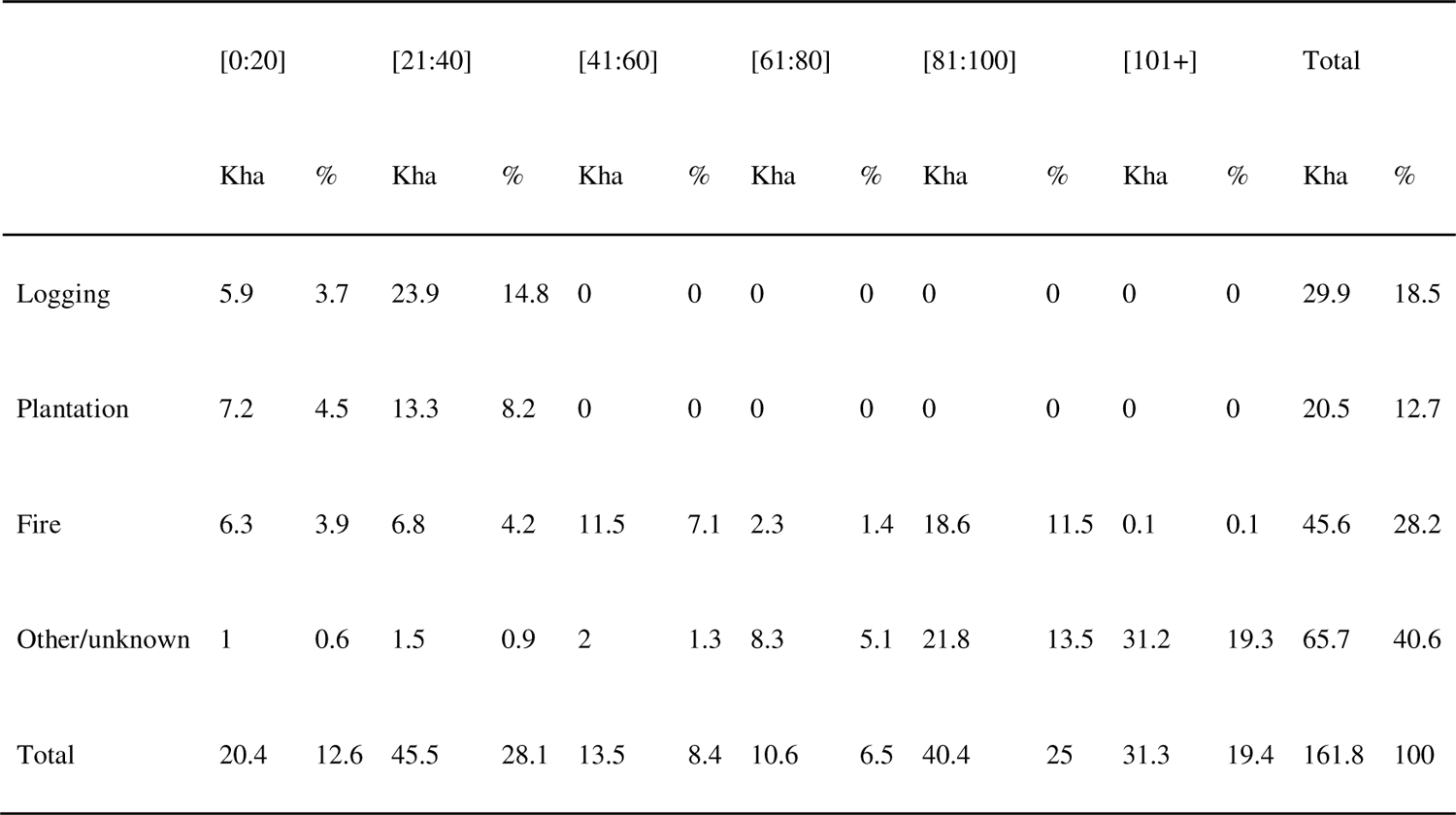
Burned areas by age class (columns) and pre-2023 disturbance types (rows) for productive forests in the **UG 102**. Results are expressed in 1000 ha (Kha) and % of the total.

|  | [0:20] |  | [21:40] |  | [41:60] |  | [61:80] |  | [81:100] |  | [101+] |  | Total |  |
| --- | --- | --- | --- | --- | --- | --- | --- | --- | --- | --- | --- | --- | --- | --- |
|  | Kha | % | Kha | % | Kha | % | Kha | % | Kha | % | Kha | % | Kha | % |
| Logging | 5.9 | 3.7 | 23.9 | 14.8 | 0 | 0 | 0 | 0 | 0 | 0 | 0 | 0 | 29.9 | 18.5 |
| Plantation | 7.2 | 4.5 | 13.3 | 8.2 | 0 | 0 | 0 | 0 | 0 | 0 | 0 | 0 | 20.5 | 12.7 |
| Fire | 6.3 | 3.9 | 6.8 | 4.2 | 11.5 | 7.1 | 2.3 | 1.4 | 18.6 | 11.5 | 0.1 | 0.1 | 45.6 | 28.2 |
| Other/unknown | 1 | 0.6 | 1.5 | 0.9 | 2 | 1.3 | 8.3 | 5.1 | 21.8 | 13.5 | 31.2 | 19.3 | 65.7 | 40.6 |
| Total | 20.4 | 12.6 | 45.5 | 28.1 | 13.5 | 8.4 | 10.6 | 6.5 | 40.4 | 25 | 31.3 | 19.4 | 161.8 | 100 |

**Table S4.11.** Burned areas by age class (columns) and composition groups (rows) for productive forests in the **UG 042**. Results are expressed in 1000 ha (Kha) and % of the total.

|  | [0:20] |  | [21:40] |  | [41:60] |  | [61:80] |  | [81:100] |  | [101+] |  | Total |  |
| --- | --- | --- | --- | --- | --- | --- | --- | --- | --- | --- | --- | --- | --- | --- |
|  | Kha | % | Kha | % | Kha | % | Kha | % | Kha | % | Kha | % | Kha | % |
| Black spruce | 0.7 | 3.3 | 0.1 | 0.6 | 0.8 | 3.7 | 1.8 | 8.5 | 1.4 | 6.6 | 6.7 | 31.1 | 11.6 | 53.7 |
| Jack pine-Black sp. | 0 | 0.2 | 0 | 0.1 | 0.1 | 0.4 | 0.4 | 1.8 | 0.6 | 2.8 | 0.3 | 1.2 | 1.4 | 6.5 |
| Conifers (oth/undif) | 2.5 | 11.7 | 1.4 | 6.7 | 0 | 0.2 | 0.1 | 0.4 | 0.1 | 0.3 | 0.4 | 2 | 4.6 | 21.2 |
| Mixed-hardwood | 1.9 | 8.8 | 0.3 | 1.5 | 0.1 | 0.6 | 0.6 | 2.9 | 0.1 | 0.5 | 0.3 | 1.3 | 3.3 | 15.6 |
| Unknown | 0.6 | 2.9 | 0 | 0.2 | 0 | 0 | 0 | 0 | 0 | 0 | 0 | 0 | 0.7 | 3.1 |
| Total | 5.8 | 27 | 1.9 | 9 | 1 | 4.8 | 2.9 | 13.6 | 2.2 | 10.1 | 7.6 | 35.5 | 21.5 | 100 |

**Table S4.12.** Burned areas by age class (columns) and pre-2023 disturbance types (rows) for productive forests in the **UG 042**. Results are expressed in 1000 ha (Kha) and % of the total.

|  | [0:20] |  | [21:40] |  | [41:60] |  | [61:80] |  | [81:100] |  | [101+] |  | Total |  |
| --- | --- | --- | --- | --- | --- | --- | --- | --- | --- | --- | --- | --- | --- | --- |
|  | Kha | % | Kha | % | Kha | % | Kha | % | Kha | % | Kha | % | Kha | % |
| Logging | 3.5 | 16.4 | 1.6 | 7.3 | 0 | 0 | 0 | 0 | 0 | 0 | 0 | 0 | 5.1 | 23.8 |
| Plantation | 0.8 | 3.9 | 0.2 | 0.8 | 0 | 0 | 0 | 0 | 0 | 0 | 0 | 0 | 1 | 4.7 |
| Fire | 0 | 0 | 0 | 0 | 0 | 0 | 0.6 | 2.7 | 0.5 | 2.3 | 0.1 | 0.6 | 1.2 | 5.5 |
| Other/unknown | 1.4 | 6.7 | 0.2 | 0.8 | 1 | 4.8 | 2.4 | 11 | 1.7 | 7.8 | 7.5 | 34.9 | 14.2 | 66 |
| Total | 5.8 | 27 | 1.9 | 9 | 1 | 4.8 | 2.9 | 13.6 | 2.2 | 10.1 | 7.6 | 35.5 | 21.5 | 100 |

**Table S4.13.** Burned areas by age class (columns) and composition groups (rows) for productive forests in the **UG 025**. Results are expressed in 1000 ha (Kha) and % of the total.

|  | [0:20] |  | [21:40] |  | [41:60] |  | [61:80] |  | [81:100] |  | [101+] |  | Total |  |
| --- | --- | --- | --- | --- | --- | --- | --- | --- | --- | --- | --- | --- | --- | --- |
|  | Kha | % | Kha | % | Kha | % | Kha | % | Kha | % | Kha | % | Kha | % |
| Black spruce | 3.7 | 6.4 | 4 | 6.9 | 0.8 | 1.4 | 5.4 | 9.1 | 0.7 | 1.2 | 14 | 23.9 | 28.6 | 48.8 |
| Jack pine-Black sp. | 0.8 | 1.3 | 0.1 | 0.3 | 1.5 | 2.5 | 4.3 | 7.3 | 1.1 | 1.9 | 0.5 | 0.8 | 8.3 | 14.1 |
| Conifers (oth/undif) | 0.8 | 1.3 | 1.9 | 3.2 | 0.1 | 0.1 | 0.1 | 0.1 | 0 | 0.1 | 0.6 | 1 | 3.4 | 5.8 |
| Mixed-hardwood | 0.1 | 0.1 | 1.4 | 2.4 | 0 | 0 | 0.3 | 0.5 | 0 | 0 | 0.2 | 0.4 | 2.1 | 3.5 |
| Unknown | 9.1 | 15.5 | 7.1 | 12.2 | 0.1 | 0.1 | 0 | 0 | 0 | 0 | 0 | 0 | 16.3 | 27.8 |
| Total | 14.4 | 24.6 | 14.6 | 24.9 | 2.4 | 4.1 | 10 | 17.1 | 1.9 | 3.2 | 15.2 | 26 | 58.6 | 100 |

**Table S4.14.** Burned areas by age class (columns) and pre-2023 disturbance types (rows) for productive forests in the **UG 025**. Results are expressed in 1000 ha (Kha) and % of the total.

|  | [0:20] |  | [21:40] |  | [41:60] |  | [61:80] |  | [81:100] |  | [101+] |  | Total |  |
| --- | --- | --- | --- | --- | --- | --- | --- | --- | --- | --- | --- | --- | --- | --- |
|  | Kha | % | Kha | % | Kha | % | Kha | % | Kha | % | Kha | % | Kha | % |
| Logging | 9.6 | 16.3 | 9 | 15.4 | 0 | 0 | 0 | 0 | 0 | 0 | 0 | 0 | 18.6 | 31.7 |
| Plantation | 4.4 | 7.5 | 4 | 6.8 | 0 | 0 | 0 | 0 | 0 | 0 | 0 | 0 | 8.4 | 14.3 |
| Fire | 0.1 | 0.2 | 1.2 | 2.1 | 0.2 | 0.4 | 0.2 | 0.4 | 0.1 | 0.1 | 0 | 0 | 1.8 | 3.1 |
| Other/unknown | 0.3 | 0.6 | 0.4 | 0.7 | 2.2 | 3.7 | 9.8 | 16.8 | 1.8 | 3.1 | 15.2 | 26 | 29.8 | 50.9 |
| Total | 14.4 | 24.6 | 14.6 | 24.9 | 2.4 | 4.1 | 10 | 17.1 | 1.9 | 3.2 | 15.2 | 26 | 58.6 | 100 |

**Table S4.15.** Burned areas by age class (columns) and composition groups (rows) for productive forests in the **UG 027**. Results are expressed in 1000 ha (Kha) and % of the total.

|  | [0:20] |  | [21:40] |  | [41:60] |  | [61:80] |  | [81:100] |  | [101+] |  | Total |  |
| --- | --- | --- | --- | --- | --- | --- | --- | --- | --- | --- | --- | --- | --- | --- |
|  | Kha | % | Kha | % | Kha | % | Kha | % | Kha | % | Kha | % | Kha | % |
| Black spruce | 0.1 | 0.6 | 1.7 | 13.9 | 1.7 | 13.8 | 0.2 | 1.9 | 0.6 | 4.6 | 0.8 | 6.8 | 5 | 41.5 |
| Jack pine-Black sp. | 0 | 0.2 | 0.2 | 2 | 0.3 | 2.7 | 0.3 | 2.3 | 0 | 0.3 | 0 | 0.1 | 0.9 | 7.6 |
| Conifers (oth/undif) | 0.1 | 0.6 | 2.4 | 19.7 | 0.9 | 7.3 | 0.1 | 0.9 | 0.1 | 1.1 | 0.2 | 1.4 | 3.8 | 31.1 |
| Mixed-hardwood | 0 | 0.2 | 0.5 | 4.3 | 0.9 | 7.6 | 0 | 0.3 | 0.1 | 0.6 | 0 | 0.3 | 1.6 | 13.3 |
| Unknown | 0.6 | 5 | 0.1 | 0.8 | 0.1 | 0.5 | 0 | 0.2 | 0 | 0 | 0 | 0 | 0.8 | 6.5 |
| Total | 0.8 | 6.6 | 4.9 | 40.7 | 3.9 | 31.9 | 0.7 | 5.6 | 0.8 | 6.6 | 1 | 8.6 | 12.1 | 100 |

**Table S4.16.**
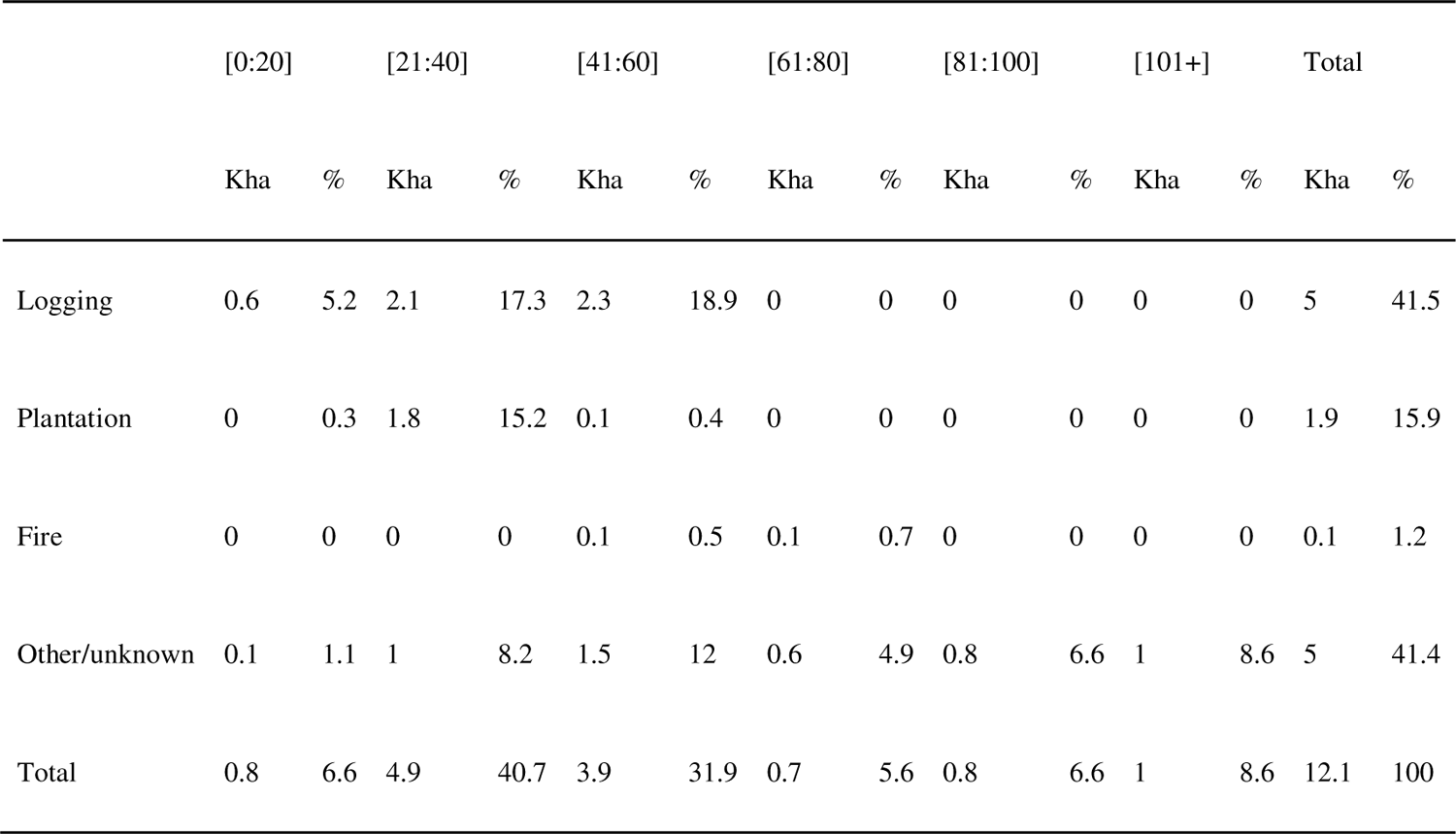
Burned areas by age class (columns) and pre-2023 disturbance types (rows) for productive forests in the **UG 027**. Results are expressed in 1000 ha (Kha) and % of the total.

|  | [0:20] |  | [21:40] |  | [41:60] |  | [61:80] |  | [81:100] |  | [101+] |  | Total |  |
| --- | --- | --- | --- | --- | --- | --- | --- | --- | --- | --- | --- | --- | --- | --- |
|  | Kha | % | Kha | % | Kha | % | Kha | % | Kha | % | Kha | % | Kha | % |
| Logging | 0.6 | 5.2 | 2.1 | 17.3 | 2.3 | 18.9 | 0 | 0 | 0 | 0 | 0 | 0 | 5 | 41.5 |
| Plantation | 0 | 0.3 | 1.8 | 15.2 | 0.1 | 0.4 | 0 | 0 | 0 | 0 | 0 | 0 | 1.9 | 15.9 |
| Fire | 0 | 0 | 0 | 0 | 0.1 | 0.5 | 0.1 | 0.7 | 0 | 0 | 0 | 0 | 0.1 | 1.2 |
| Other/unknown | 0.1 | 1.1 | 1 | 8.2 | 1.5 | 12 | 0.6 | 4.9 | 0.8 | 6.6 | 1 | 8.6 | 5 | 41.4 |
| Total | 0.8 | 6.6 | 4.9 | 40.7 | 3.9 | 31.9 | 0.7 | 5.6 | 0.8 | 6.6 | 1 | 8.6 | 12.1 | 100 |

**Table S4.17.** Burned areas by age class (columns) and composition groups (rows) for productive forests in the **UG 024**. Results are expressed in 1000 ha (Kha) and % of the total.

|  | [0:20] |  | [21:40] |  | [41:60] |  | [61:80] |  | [81:100] |  | [101+] |  | Total |  |
| --- | --- | --- | --- | --- | --- | --- | --- | --- | --- | --- | --- | --- | --- | --- |
|  | Kha | % | Kha | % | Kha | % | Kha | % | Kha | % | Kha | % | Kha | % |
| Black spruce | 0.1 | 1.4 | 0 | 0.2 | 0 | 0.4 | 0.2 | 5.3 | 0.8 | 18.9 | 0.9 | 20.1 | 2 | 46.4 |
| Jack pine-Black sp. | 0.1 | 1.6 | 0 | 0 | 0 | 0 | 0 | 0.9 | 1 | 24.1 | 0 | 0.2 | 1.1 | 26.7 |
| Conifers (oth/undif) | 0.2 | 5.2 | 0.1 | 2.1 | 0 | 0 | 0 | 0.5 | 0 | 0 | 0 | 0.8 | 0.4 | 8.8 |
| Mixed-hardwood | 0.1 | 3.5 | 0 | 1 | 0 | 0 | 0 | 0.1 | 0.1 | 1.5 | 0 | 0.1 | 0.3 | 6.2 |
| Unknown | 0.4 | 10.4 | 0.1 | 1.4 | 0 | 0 | 0 | 0 | 0 | 0 | 0 | 0 | 0.5 | 11.8 |
| Total | 0.9 | 22.2 | 0.2 | 4.8 | 0 | 0.4 | 0.3 | 6.9 | 1.9 | 44.5 | 0.9 | 21.2 | 4.2 | 100 |

**Table S4.18.**
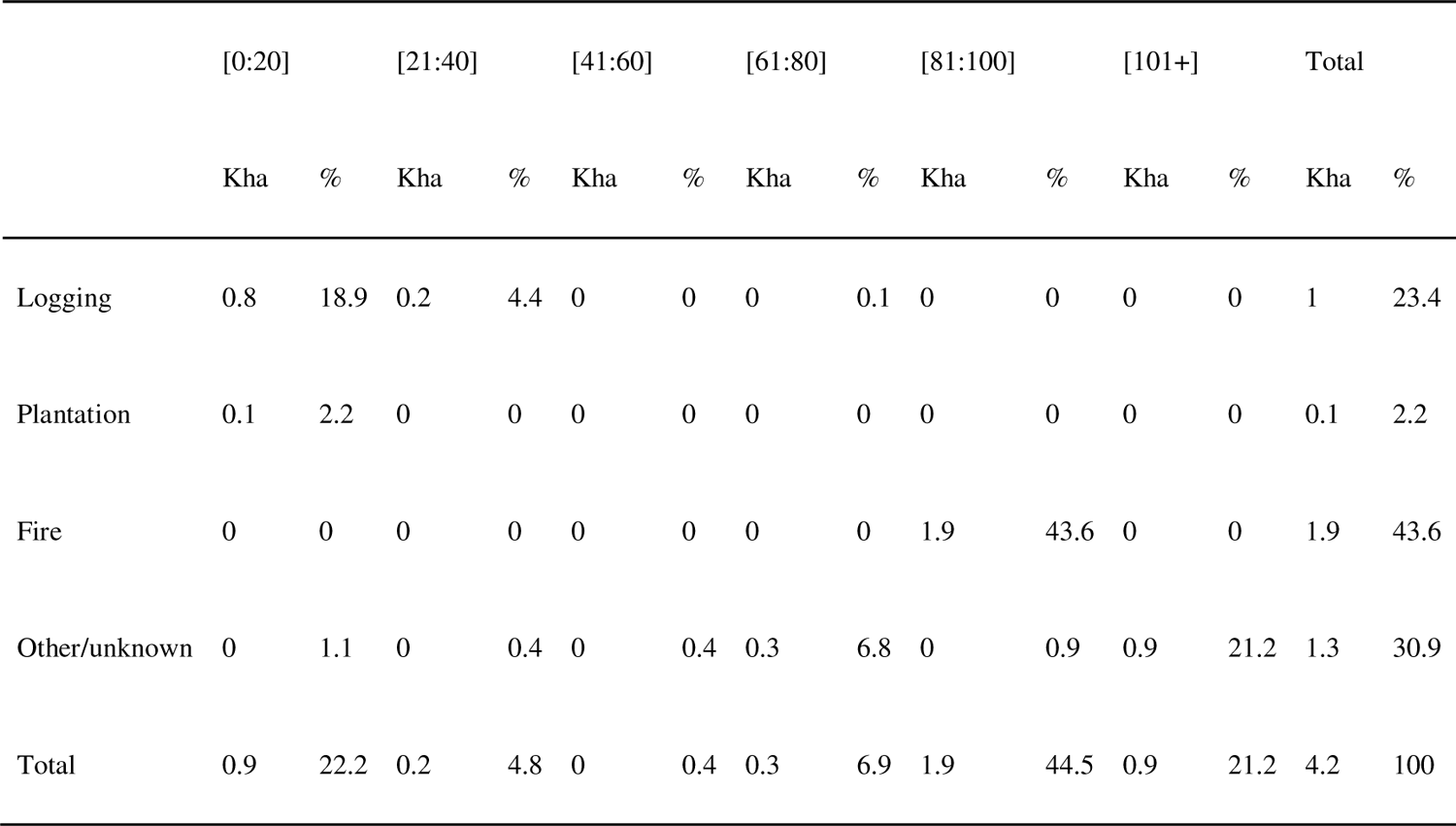
Burned areas by age class (columns) and pre-2023 disturbance types (rows) for productive forests in the **UG 024**. Results are expressed in 1000 ha (Kha) and % of the total.

**Table S4.19.** Burned areas by age class (columns) and composition groups (rows) for productive forests in the **UG 097**. Results are expressed in 1000 ha (Kha) and % of the total.

|  | [0:20] |  | [21:40] |  | [41:60] |  | [61:80] |  | [81:100] |  | [101+] |  | Total |  |
| --- | --- | --- | --- | --- | --- | --- | --- | --- | --- | --- | --- | --- | --- | --- |
|  | Kha | % | Kha | % | Kha | % | Kha | % | Kha | % | Kha | % | Kha | % |
| Black spruce | 0.7 | 3.3 | 0.2 | 1 | 0.1 | 0.5 | 0.8 | 3.9 | 0.9 | 4.3 | 5.9 | 27.8 | 8.7 | 40.8 |
| Jack pine-Black sp. | 0.1 | 0.4 | 0 | 0 | 0 | 0 | 0.1 | 0.4 | 0.6 | 3 | 0.6 | 2.8 | 1.4 | 6.6 |
| Conifers (oth/undif) | 0.7 | 3.5 | 2.4 | 11.4 | 0.1 | 0.5 | 0.1 | 0.4 | 0 | 0.2 | 2.3 | 10.6 | 5.6 | 26.5 |
| Mixed-hardwood | 0.3 | 1.2 | 1.1 | 5 | 0 | 0.2 | 0.1 | 0.4 | 0 | 0.1 | 0 | 0.2 | 1.5 | 7 |
| Unknown | 2.6 | 12.3 | 1.3 | 6 | 0.1 | 0.7 | 0 | 0 | 0 | 0 | 0 | 0 | 4 | 19 |
| Total | 4.4 | 20.6 | 5 | 23.4 | 0.4 | 1.9 | 1.1 | 5.1 | 1.6 | 7.6 | 8.8 | 41.5 | 21.3 | 100 |

**Table S4.20.** Burned areas by age class (columns) and pre-2023 disturbance types (rows) for productive forests in the **UG 097**. Results are expressed in 1000 ha (Kha) and % of the total.

|  | [0:20] |  | [21:40] |  | [41:60] |  | [61:80] |  | [81:100] |  | [101+] |  | Total |  |
| --- | --- | --- | --- | --- | --- | --- | --- | --- | --- | --- | --- | --- | --- | --- |
|  | Kha | % | Kha | % | Kha | % | Kha | % | Kha | % | Kha | % | Kha | % |
| Logging | 3.3 | 15.5 | 4.5 | 21.4 | 0 | 0 | 0 | 0 | 0 | 0 | 0 | 0 | 7.9 | 36.9 |
| Plantation | 0.5 | 2.3 | 0 | 0 | 0 | 0 | 0 | 0 | 0 | 0 | 0 | 0 | 0.5 | 2.4 |
| Fire | 0 | 0.1 | 0.2 | 0.8 | 0.2 | 1.2 | 0.1 | 0.5 | 0 | 0 | 0 | 0 | 0.5 | 2.5 |
| Other/unknown | 0.6 | 2.7 | 0.3 | 1.2 | 0.1 | 0.7 | 1 | 4.6 | 1.6 | 7.6 | 8.8 | 41.5 | 12.4 | 58.2 |
| Total | 4.4 | 20.6 | 5 | 23.4 | 0.4 | 1.9 | 1.1 | 5.1 | 1.6 | 7.6 | 8.8 | 41.5 | 21.3 | 100 |

**Table S4.21.** Burned areas by age class (columns) and composition groups (rows) for productive forests in the **UG 093**. Results are expressed in 1000 ha (Kha) and % of the total.

|  | [0:20] |  | [21:40] |  | [41:60] |  | [61:80] |  | [81:100] |  | [101+] |  | Total |  |
| --- | --- | --- | --- | --- | --- | --- | --- | --- | --- | --- | --- | --- | --- | --- |
|  | Kha | % | Kha | % | Kha | % | Kha | % | Kha | % | Kha | % | Kha | % |
| Black spruce | 0.1 | 1.3 | 0.1 | 0.8 | 0.4 | 4.4 | 2.6 | 29.2 | 0.2 | 1.9 | 1.6 | 17.4 | 5 | 55.1 |
| Jack pine-Black sp. | 0 | 0 | 0 | 0 | 0 | 0 | 0.3 | 2.8 | 0.1 | 0.6 | 0 | 0 | 0.3 | 3.4 |
| Conifers (oth/undif) | 0 | 0.5 | 0.1 | 1 | 0.3 | 3.8 | 0.5 | 5.7 | 0 | 0.1 | 1.3 | 14.2 | 2.3 | 25.3 |
| Mixed-hardwood | 0.1 | 0.9 | 0.1 | 1.2 | 0.4 | 4.9 | 0.3 | 3.7 | 0 | 0.2 | 0.2 | 2.6 | 1.2 | 13.4 |
| Unknown | 0.2 | 2.6 | 0 | 0 | 0 | 0 | 0 | 0.1 | 0 | 0 | 0 | 0 | 0.2 | 2.7 |
| Total | 0.5 | 5.4 | 0.3 | 3 | 1.2 | 13.1 | 3.8 | 41.5 | 0.3 | 2.8 | 3.1 | 34.2 | 9 | 100 |

**Table S4.22.** Burned areas by age class (columns) and pre-2023 disturbance types (rows) for productive forests in the **UG 093**. Results are expressed in 1000 ha (Kha) and % of the total.

|  | [0:20] |  | [21:40] |  | [41:60] |  | [61:80] |  | [81:100] |  | [101+] |  | Total |  |
| --- | --- | --- | --- | --- | --- | --- | --- | --- | --- | --- | --- | --- | --- | --- |
|  | Kha | % | Kha | % | Kha | % | Kha | % | Kha | % | Kha | % | Kha | % |
| Logging | 0.1 | 0.6 | 0.2 | 2.2 | 1.1 | 11.7 | 3.4 | 38 | 0 | 0 | 0 | 0 | 4.7 | 52.4 |
| Plantation | 0 | 0 | 0 | 0 | 0 | 0 | 0 | 0 | 0 | 0 | 0 | 0 | 0 | 0 |
| Fire | 0 | 0.4 | 0 | 0 | 0 | 0 | 0.3 | 3.4 | 0 | 0 | 0 | 0 | 0.3 | 3.8 |
| Other/unknown | 0.4 | 4.4 | 0.1 | 0.8 | 0.1 | 1.4 | 0 | 0.2 | 0.3 | 2.8 | 3.1 | 34.2 | 4 | 43.8 |
| Total | 0.5 | 5.4 | 0.3 | 3 | 1.2 | 13.1 | 3.8 | 41.5 | 0.3 | 2.8 | 3.1 | 34.2 | 9 | 100 |

**Figure S4.2.**
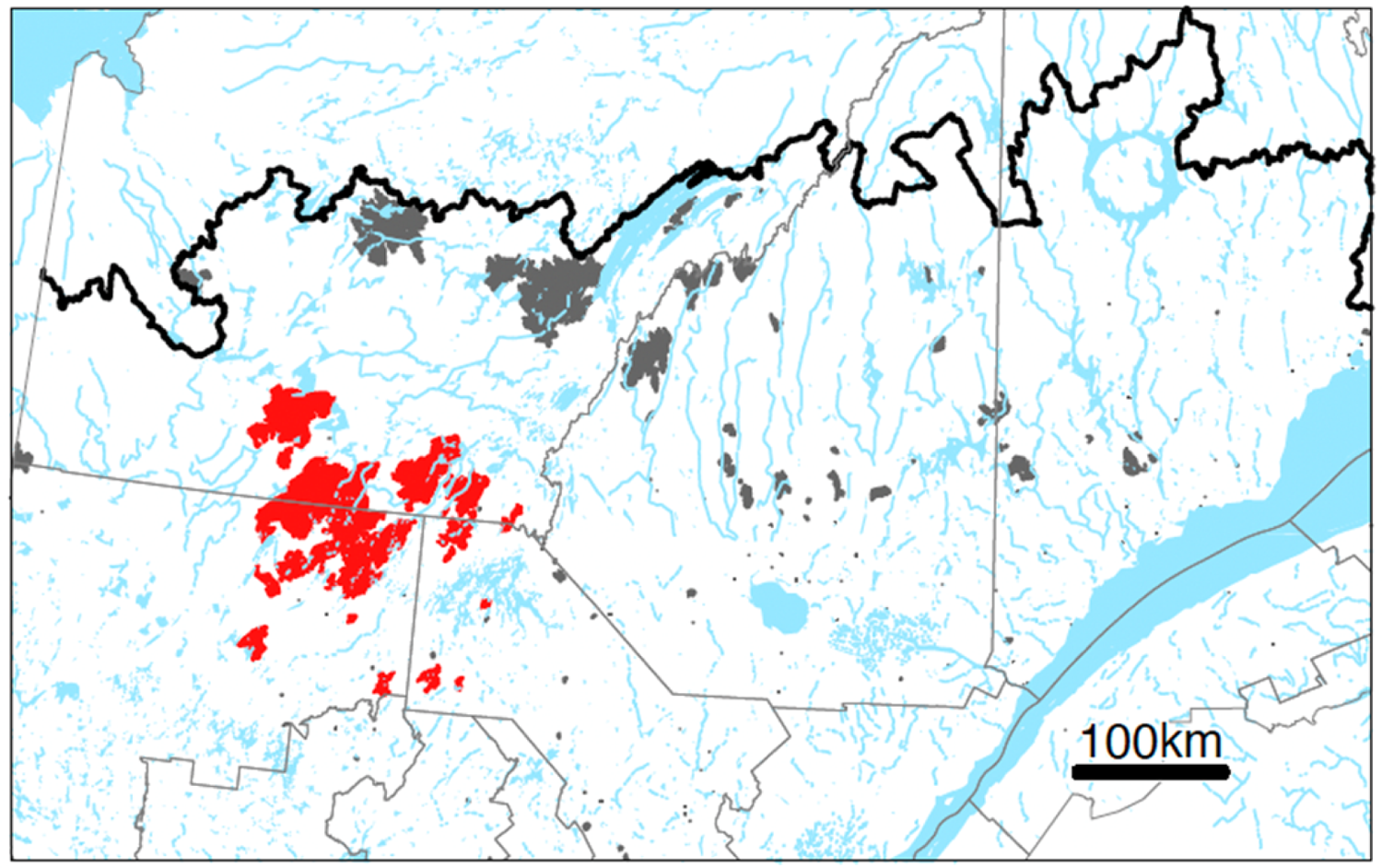
Landscapes retained (in red) for the fire selectivity analysis (Figures S3 to S5). Other landscapes affected by fires are shown in dark gray. The thick black line in the upper part of the map shows the northern limit of managed forests, and fine grey lines show the boundaries of Québec’s administrative region.

**Figure S4.3.**
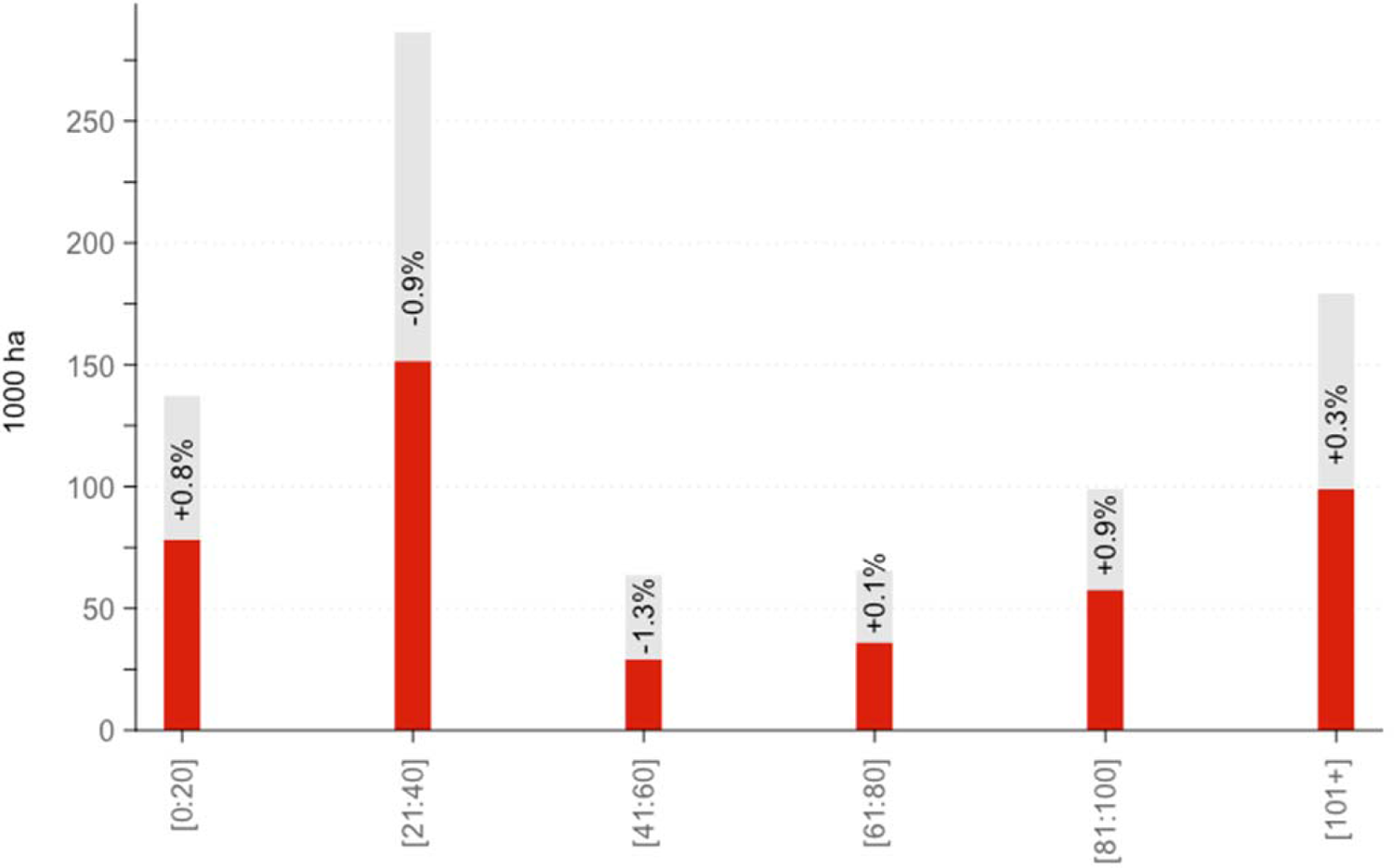
Differences between areas burned by age class and their availability in the landscape. For each age class, the top of the red bar indicates the total area burned, and the top of the gray bar indicates its availability in the landscape. The % values indicate the difference between an age class proportion of total burn area and its proportion of total abundance within the landscape (i.e., % burned minus % available).

**Figure S4.4.**
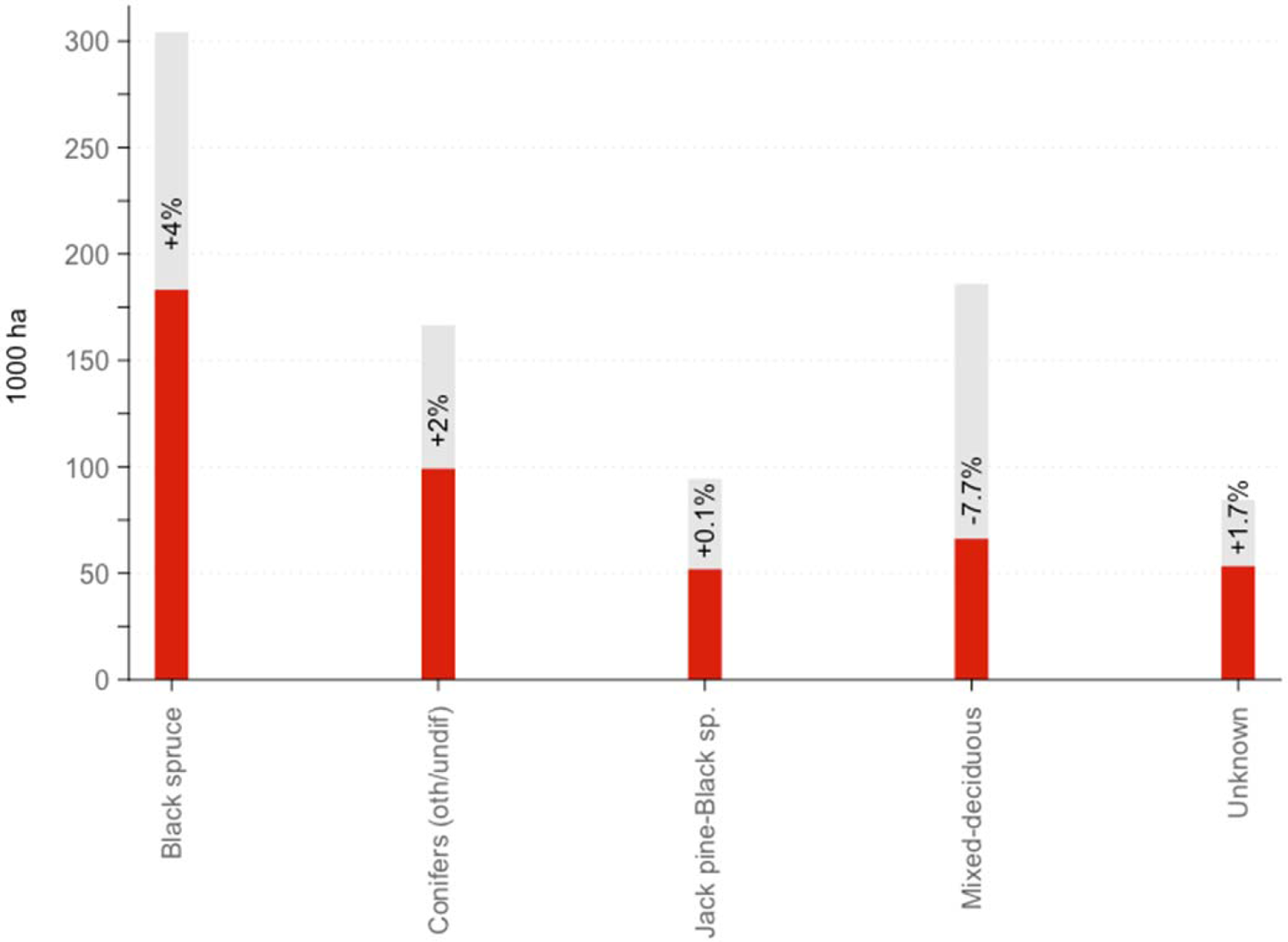
Differences between areas burned by composition groups and their availability in the landscape. For each composition group, the top of the red bar indicates the total area burned, and the top of the gray bar indicates its availability in the landscape. The % values indicate the difference between a composition group’s proportion of total burn area and its proportion of total abundance within the landscape (i.e., % burned minus % available).

**Figure S4.5.**
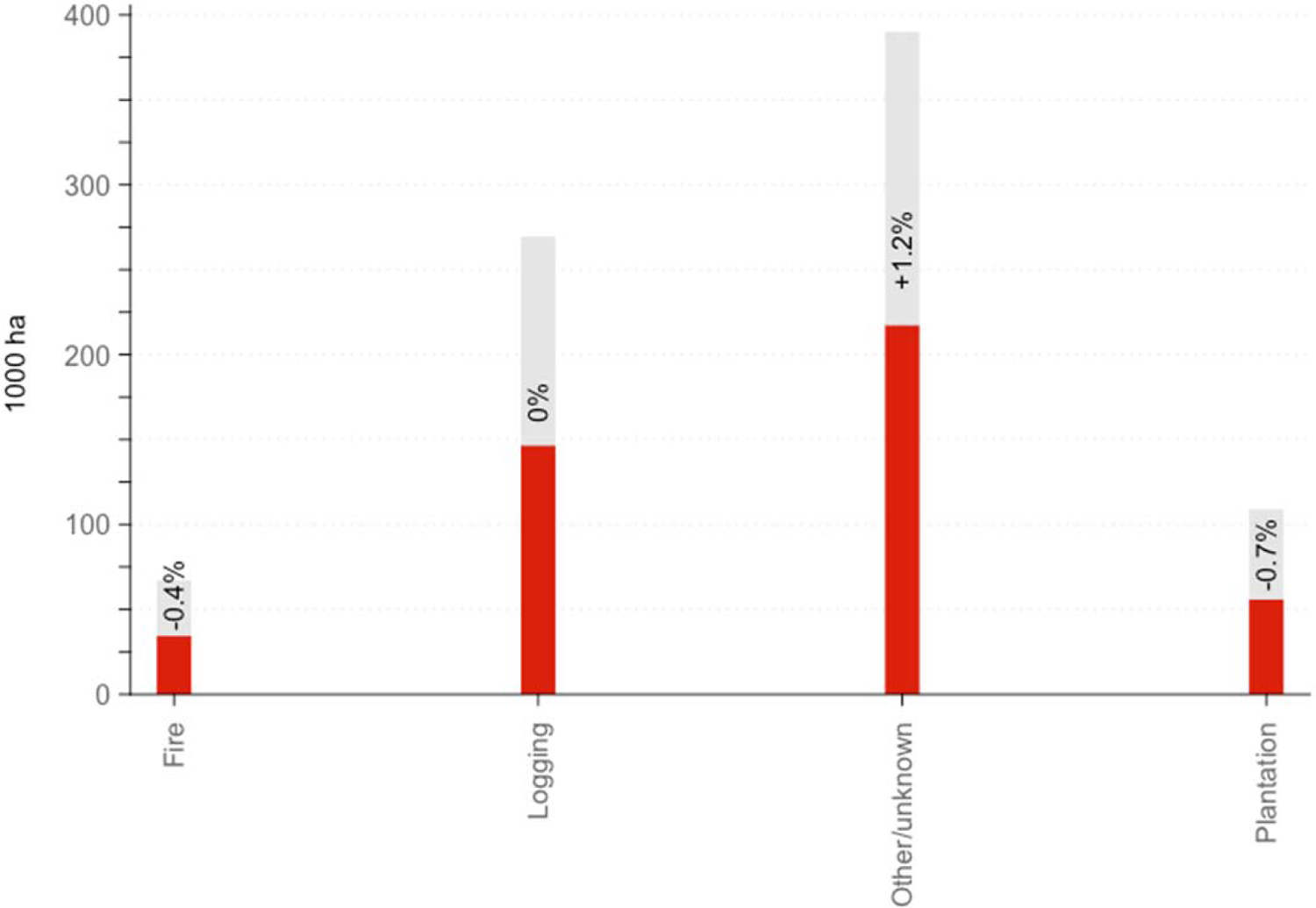
Differences between areas burned by pre-2023 disturbance types and their availability in the landscape. For each pre-2023 disturbance type, the top of the red bar indicate the total area burned, and the top of the gray bar indicates its availability in the landscape. The % values indicate the difference between a pre-2023 disturbance-type proportion of total burn area and its proportion of total abundance within the landscape (i.e., % burned minus % available).

